# Epigenomic manipulation reveals the relationship between locus specific chromatin dynamics and gene expression

**DOI:** 10.1101/2024.07.17.603961

**Authors:** Vrinda Venu, Eric M. Small, Cullen Roth, Samantha H. Adikari, Anna Hendrika Cornelia Vlot, Kyle A. Sullivan, Chanaka Roshan Abeyratne, Daniel Jacobson, Shawn R. Starkenburg, Karissa Y. Sanbonmatsu, Christina R. Steadman

## Abstract

Dysregulation of epigenetic processes leads to a plethora of abnormalities including disease states such as cancer. Therapies focused on epigenetic modulation alter gene expression to correct dysfunction, though the mechanisms and perpetuation of these states is unknown. Here, we use integrated epigenomics and three-dimensional chromatin structure-function analyses after acute histone deacetylase inhibitor cancer drug treatment (suberoylanilide hydroxamic acid in lung cancer cells). Treatment induced substantial (13%) genomic rearrangement that rebounds despite persistent gene expression changes and spreading of acetylation. The chromatin functional landscape (accessibility, active transcription modification, and gene expression) is controlled and locus-specific, while chromatin contacts are globally altered resulting in a moderate weakening of topologically associating domains. Chromatin states are more dynamic at transcriptionally active loci while genes with reduced expression are epigenetically stable suggesting chromatin architectural turnover and nucleosome remodeling is locus-specific and underlies the bidirectional expression changes. Thus, local 3D chromatin and genome structural dynamics is integral for loci regulation in response to epigenomic perturbation. The partial persistence of these altered features may have larger implications for efficacy of epigenetic drugs in amelioration of disease states.

## Introduction

Epigenetic processes constitute a series of regulatory mechanisms that cells employ to integrate and process external stimuli, ultimately resulting in the support and control of genomic function without altering the underlying sequence. This fundamental layer of regulation is manifested as chemical modification of chromatin components (histones and DNA), which are thought to alter accessibility of transcriptional machinery, thereby imparting significant influence on gene expression and ultimately, phenotype. Dysregulation of integral epigenetic processes leads to a plethora of abnormalities, ranging from developmental aberrations to disease states such as cancer. As such, direct targeting of modifications using epigenetic therapies focuses on controlling gene expression to correct dysregulation without persistence (Baylin, 2005; Morgan and Shilatifard, 2015; Zahir and Brown, 2011), though in many cases, the level of permanency has not been assessed. Epigenetic therapies utilize drugs that inhibit the activity or binding of chromatin remodeling enzymes that are responsible for imparting, removing, or remodeling epigenetic modifications or chromatin features (Keppler and Archer, 2008). In particular, histone deacetylase inhibitors (HDACi) block enzymes that remove histone acetylation, resulting in either substantially increased acetylation or retention of acetylation that would have otherwise been removed. For its part, acetylation is thought to preferentially associate (or induce) open chromatin regions, thereby allowing for transcription factor binding and eventual gene expression. HDACi therapies are currently used in the clinical setting to treat a variety of diseases ranging from cancer (Li and Seto, 2016) to neurobiological disorders (Chuang et al., 2009). The efficacy of HDACi treatment has been widely explored particularly in relation to the progression of cancer, where treatment reduces tumor cell growth while concurrently increasing the expression of tumor suppressor genes and decreasing the expression of oncogenes (Glozak and Seto, 2007; Halsall et al., 2015; Sanchez et al., 2018). Interestingly, reduced (hypo)acetylation of histones induced by epigenetic therapies that inhibit histone acetyltransferase enzymes also produce similar anti-tumor functional outcomes (Hogg et al., 2021). Potentially, perturbation of acetylation homeostasis triggers a non-proliferative transcriptional program during treatment. Despite these observations, how bidirectional gene expression occurs in response to these global changes in acetylation is not well understood. Deeper knowledge of these processes is warranted given the presumed role of acetylation, its correlation to actively transcribed genes in non-disease states, the use of HDACi in clinical settings, and the persistence of drug resistance even for efficacious treatments.

The intricate interplay between epigenetic modifications and gene expression potentially impacts—or is mediated by—genomic structures like compartmentalization for regulation of entire genomic loci. In short, the functionality of the genome is intimately intertwined with its structural features and organization in three-dimensional (3D) space, including chromatin contact frequency, distribution, loop formation, and phase separation (Finn et al., 2019; Hnisz et al., 2017; Misteli, 2020). Indeed, the impact of environmental perturbations, including those that contribute to the development and progression of disease states, is mediated by dynamic chromatin remodeling in addition to epigenetic changes. These local features are potentially connected to larger global chromatin structures (Glozak and Seto, 2007; Wolffe, 2001). The development and utilization of genome-wide chromatin capture technologies, such as Hi-C, delineate chromosomal rearrangements, annotation of genomic events, and global pictures of 3D spatial configurations of chromatin inside the nucleus. These analyses, combined with nucleosome occupancy from ATAC-seq assays and epigenetic modification measurements from ChIP-seq experiments can provide visualization of the entire chromatin landscape at various resolutions, ultimately allowing for deeper knowledge of the dynamic interactions between epigenetic modifications and chromatin structure-function relationships (Hogg et al., 2021; Venu et al., 2023). Thus, a clear understanding of chromatin responses, including global and fine structural features, and epigenetic modifications when acetylation homeostasis is perturbed by an epigenetic therapy will help elucidate how functional outcomes occur and potentially persist.

Here, we present a comprehensive, genome-wide investigation to ascertain the impact of HDACi treatment on global and local chromatin structure, epigenomics, and genome function. Chromatin responses were assessed in human lung carcinoma cells (A549) after treatment with suberoylanilide hydroxamic acid (SAHA) at two timepoints post treatment (24 and 48 hours). SAHA is the first FDA-approved drug specifically targeted to HDAC enzymes, with non-specific inhibition of multiple HDACs including HDAC1, HDAC2, HDAC3, and HDAC6 (Beckers et al., 2007; Li and Seto, 2016). This experimental design serves as a framework for investigating chromatin 3D structure-function dynamics over time, with the 24-hour time point providing information on cellular responses from direct interaction with the drug, while the 48-hour time point demonstrates the retention of HDACi effects post treatment. This is the first report to describe the impact of SAHA on chromatin features in a paired manner, including histone modifications, accessibility, and contact changes. This detailed paired investigation of chromatin structure-function dynamics illuminates the complexity and intricate nature of global and local chromatin responses to an acute epigenomic perturbation and its contribution to transcriptional regulation.

We find concurrent alterations in gene expression, chromatin accessibility, and 3D chromatin architecture to various extents, in response to HDACi drug treatment. Importantly, the global increase in acetylation in response to SAHA does not result in a genome-wide increase in chromatin accessibility nor the opening of higher-order chromatin architecture; rather, regulation of local chromatin features supports the bidirectional gene expression change. Further, the locus-specific turnover rate of chromatin structural components, including nucleosome remodeling and topologically associating domain dynamics influence the directionality of the gene expression change. Therefore, chromatin dynamics are significantly involved in shaping the cellular response to HDACi treatment. As such, investigation of these processes may be critical to improve the development and implementation of novel epigenomic therapies.

## Results

### Multi-omics characterization of chromatin response to SAHA treatment

We performed multi-omic assessments of human lung carcinoma (A549) cells with and without HDACi drug treatment to identify transcriptional, epigenomic, and chromatin structural responses. The appropriate dose and duration of suberoylanilide hydroxamic acid (SAHA) treatment was determined from a pilot dose-response study (Figure S1). A concentration of 10 µM SAHA was chosen for subsequent experiments as this treatment considerably altered acetylation on lysine 27 of histone 3 (H3K27ac) while still retaining viable cells at 48 hours post treatment for chromatin structure-function characterization. (Figure S1A). Time course experiments were conducted in triplicate; control (mock-treated) and treated (10 µM SAHA) A549 cells were collected at 24 and 48 hpt. At both timepoints, intact cells had marginally increased cell size and rounded cell morphology as identified via bright field and fluorescence microscopy (Figure S1B). Genome-wide assays for control and treated paired cells included ChIP-seq (H3, H3ac, H3K27ac, H3K4me3) and RNA-seq, each with three biological replicates, and Hi-C with two biological replicates (Figure 1A). ATAC-seq samples from three biological replicates per condition were generated from a separate repeated experiment. This experimental design generated multiple data sets allowing for inspection of global and local chromatin features, epigenetic modifications, and gene expression for determining concurrent chromatin structure-function dynamics in response to epigenetic drug treatment.

**Figure 1:**
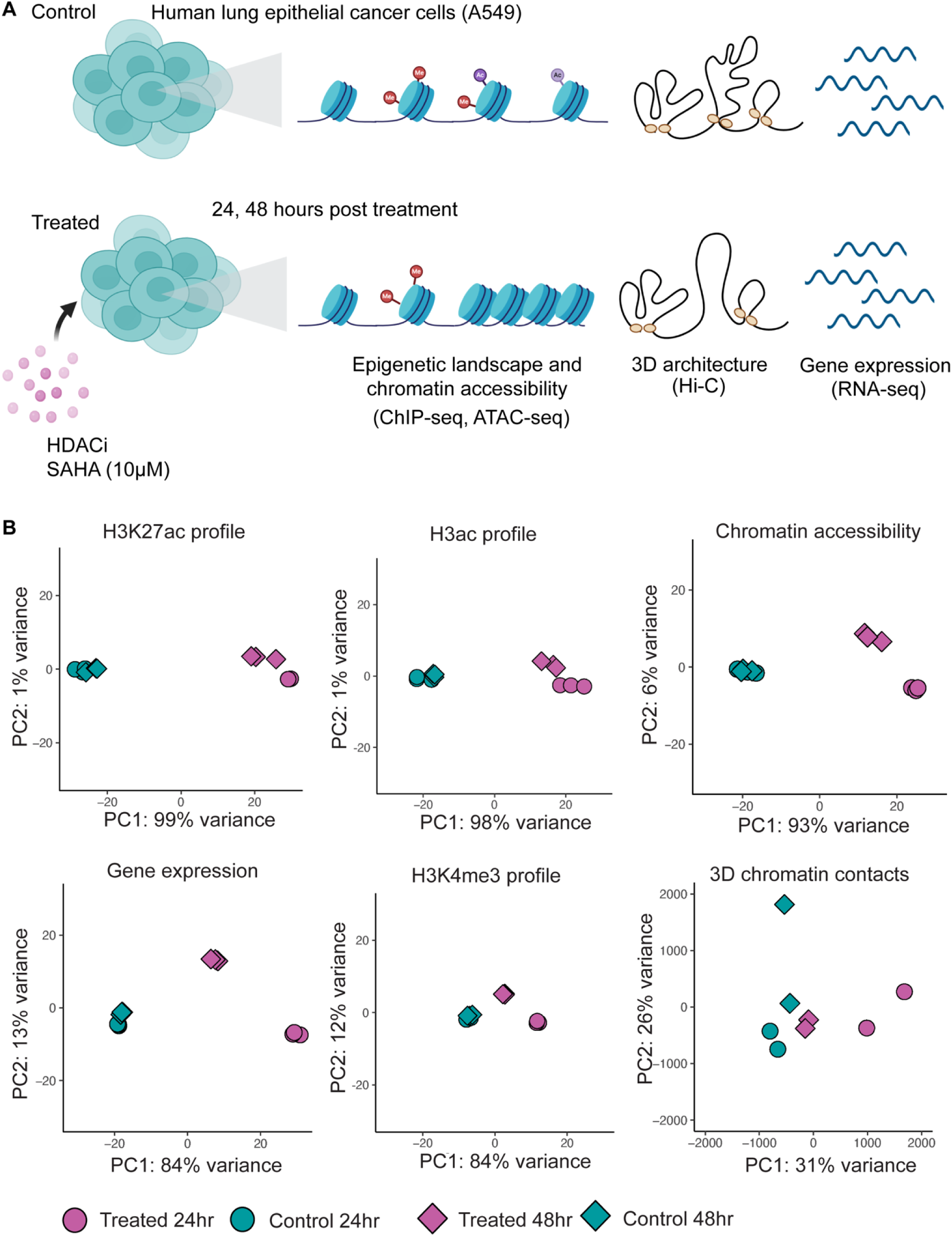
A schematic representation of the experimental design and genome-wide sample variance demonstrating the treatment effect in each data set. A) Human lung epithelial cancer cells (A549) were treated with an HDACi drug, suberoylanilide hydroxamic acid (SAHA) at 10 µM concentration. At 24 and 48 hours post treatment, Control (mock-treated) and Treated (10 µM SAHA) A549 cells from the same culture flask were divided for ChIP-seq, RNA-seq, and Hi-C experiments. ATAC-seq samples were generated from a repeated experiment. B) Principal Component Analysis (PCA) of genome-wide, histone 3 lysine 27 acetylation (H3K27ac), histone pan-acetyl (H3ac), chromatin accessibility, gene expression, histone 3 lysine 4 trimethylation (H3K4me3), and 3D chromatin contact profiles present separation based on treatment and timepoint as depicted by the first two principal components (PC1; x-axis; PC2: y-axis).

### HDAC inhibitor treatment elicits strong time-dependent responses in chromatin structure and function

Treatment with 10 µM SAHA induced substantial, significant changes in chromatin structure and function compared to mock-treated (control) cells at each timepoint as demonstrated by Principal Component Analysis (Figure 1B). While keeping high replicate concordance in all datasets (Figure S2-S5), we observed substantial separation due to SAHA treatment in the H3K27 acetylation profile (99% variance explained by PC1) and the H3ac profile (98% variance explained by PC1), which covers pan-acetylation on lysines 9, 14, 18, 23, and 27; these outcomes were expected given the direct inhibition of class I and II HDACs involved in H3K27 homeostasis (Caslini et al., 2019; Dang et al., 2022). Specific chromatin responses at 24 and 48 hpt were markedly differential for chromatin accessibility, gene expression, H3K4 trimethylation, and to some degree 3D chromatin contacts, though in all measured features, a larger difference was observed at 24 hpt. Further, while histone acetylation remains substantially different in treated cells at 48 hpt, chromatin structure and functional responses (expression and histone methylation) rebound toward control culture characteristics, though this rebound is not complete suggesting residual effects of treatment. This indicates not only a stronger effect size at 24 hpt compared to 48 hpt, potentially due to compensatory effects or reduced viability, but the resiliency of the cell line to acute stress. The variation in effect size of these different features suggests that despite global concordance, a complex non-linear relationship exists between global chromatin structural features (e.g., 3D chromatin contacts) and functional output (e.g., histone modifications and gene expression).

### The relationship between histone acetylation and chromatin accessibility is non-linear

The primary function of a histone deacetylase inhibitor, such as SAHA, is to interfere with enzymes that remove lysine acetylation in histone and non-histone proteins. As such, SAHA treatment globally increased histone acetylation levels, as reflected in the H3K27 acetylation profile and broadened hyperacetylated regions at both 24 and 48 hpt (Figure 2A, curve comparison in Figure S6A). Concurrent with the global increase in histone acetylation, we observed a moderate elevation in H3K4 trimethylation, an epigenetic modification associated with active transcription, though this effect was more pronounced at 24 than 48 hpt (Figure 2B, curve comparison in Figure S6B). Unexpectedly, there was essentially no change (a slight reduction) in global chromatin accessibility despite the elevation of epigenetic modifications associated with active transcription in response to HDACi treatment (Figure 2C, curve comparison in Figure S6C). Further, while the elevation in global acetylation remained was slightly reduced at 48 hpt compared to 24 hpt, the global H3K4 trimethylation and chromatin accessibility levels remained more similar between the two timepoints (Figure 2A-C).

**Figure 2:**
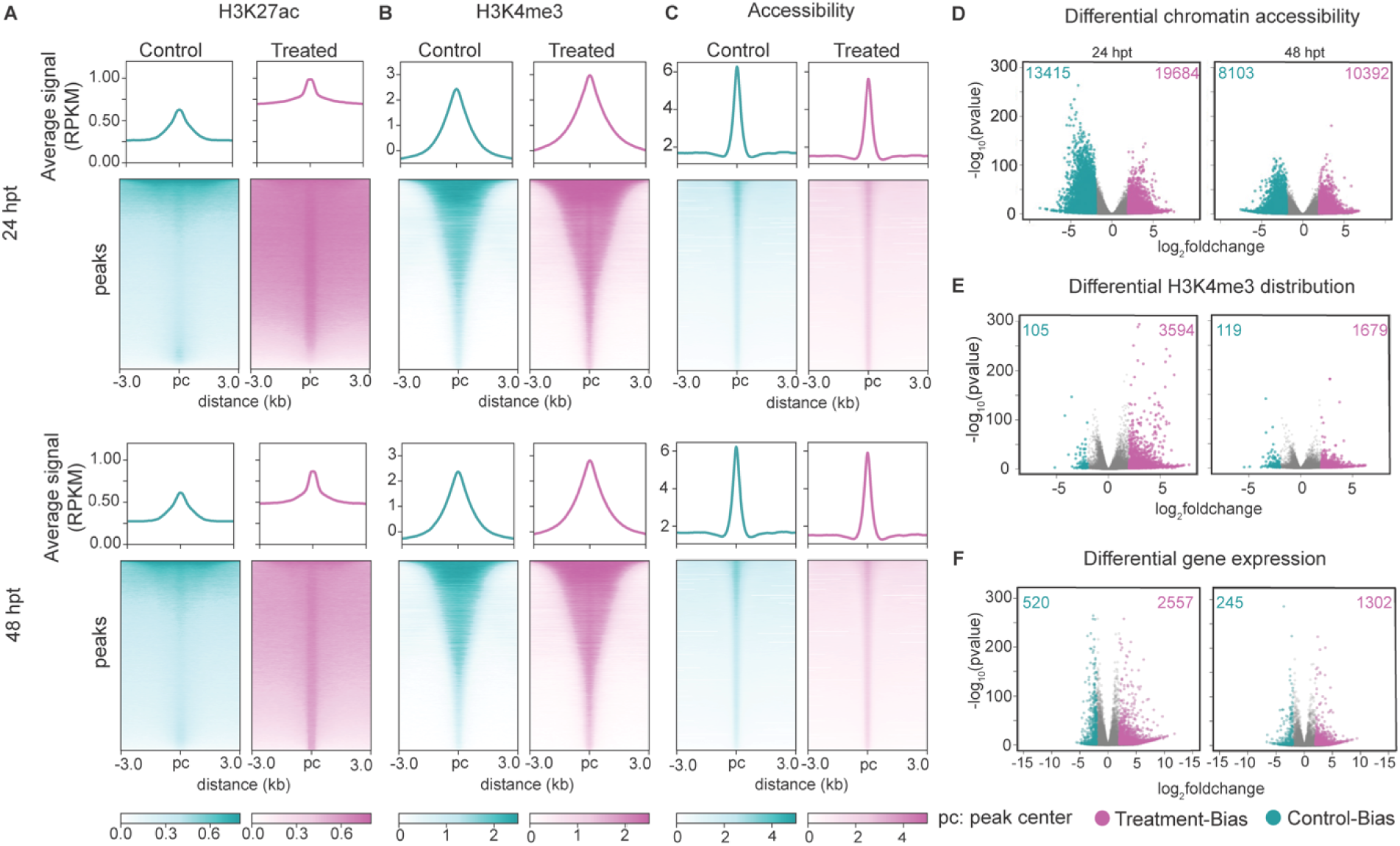
Histone acetylation (H3K27ac) and trimethylation (H3K4me3) levels are elevated without a global increase in chromatin accessibility. A) Histone acetylation (H3K27ac), B) trimethylation (H3K4me3), and C) chromatin accessibility around their respective consensus peak centers (pc) are plotted for 24 hpt and 48 hpt, where teal indicates Control (mock-treated) cells and pink indicates Treated (SAHA-treated) cells. Reads from all biological replicates are combined. The normalized average read count profile is plotted in the top panel, while the bottom panel shows the normalized read count per peak where each row represents one peak called in at least one biological replicate. Read counts were determined at ± 3 kb region around all consensus peak centers. Volcano plots demonstrate regions with differential D) chromatin accessibility, E) H3K4me3 density, and F) genes with differential expression between Control and Treated cells at 24 and 48 hpt. In all six plots, teal dots indicate significantly different (adjusted p-value <0.05) control-biased regions (log_2_ fold change <-2), where the signal is increased in control cultures (and therefore decreased in HDACi treated cultures). Pink dots indicate treatment-biased regions (adjusted P value <0.05 and log_2_ fold change >2), where the signal is increased in treated cultures (and decreased in control cultures). All remaining non-significant (adjusted p-value > 0.05) or less extreme fold changes (−2 < log2 fold change <2) are represented by grey dots.

The lack of major variation in chromatin accessibility from the global analysis warranted further investigation. When considering individual open chromatin regions (OCRs), their accessibility was both increased (pink dots) and decreased (teal dots) due to HDACi treatment (adj *p*-value < 0.05 and log 2-fold change > 0) (Figure 2D). There were more open regions with larger effects (fold change greater than two) (19,684 in Figure 2D) suggesting that HDAC inhibition led to dramatic opening of certain regions. However, there were still 13,415 regions (teal dots, Figure 2D) with decreased accessibility due to treatment. Collectively, these differentially accessible regions (both open and closed) due to treatment constitute approximately 29% and 19% of the ∼342,398 distinctly significant OCRs in the genome, and these OCRs cover approximately 5.45% of the genome. The differentially accessible OCRs are primarily associated with genic regions (∼65%) (gene body ±1kb, according to NCBI RefSeq), though 28.5% were localized to intergenic regions, and 6.5% were annotated as ‘cell type specific intergenic’ enhancers. Similarly, 25% and 18% of genes were differentially expressed (regardless of fold change) at 24 and 48 hpt respectively, whereas 64% and 46% of regions with H3K4 trimethylation had differential peaks at 24 and 48 hpt (Figure 2E, 2F). While the observed bidirectional differential gene expression has previously been reported in response to HDACi treatment (Halsall et al., 2015; Slaughter et al., 2021), we found substantially more (five times) H3K4 trimethylation peaks (Figure 2E, pink colored dots and numbers) and more upregulated genes (five times) than downregulated genes (Figure 2F, pink colored dots and numbers), both with a fold change greater than two. Collectively, these observations suggest that a major change in genome acetylation elicited rather controlled locus specific changes in chromatin functional landscape (accessibility, active transcription mark, and gene expression).

### Reduced promoter acetylation density and accessibility are associated with downregulation of gene expression potentially due to nucleosome remodeling

Next, we investigated the chromatin features that correlate with differential gene expression. Irrespective of expression status, gene promoters in treated cells possess low acetylation compared to controls (normalized to H3) with significant spreading of acetylation across gene bodies (Figure S7A). Genes upregulated (padj <0.05 and log_2_ fold change >2) by HDACi treatment possess nearly the same amount of promoter histone acetylation at 24 hpt as Controls (Figure 3A). Conversely, genes downregulated (padj <0.05 and log_2_ fold change < −2) by HDACi treatment possess considerably lower promoter acetylation 24 hpt (Figure 3B). However, nucleosome distribution is an important consideration when performing these assessments: indeed, typical normalization (to input levels) suggest decreased promoter acetylation in all differential genes (Figure S8). However, normalization to nucleosome distribution (measured via H3 ChIP-seq) as we have shown provides a more nuanced perspective of feature correlation. Thus, changes in nucleosome distribution are concurrent with differential gene expression. Further, in contrast to downregulated genes, genes upregulated after treatment possess lower promoter acetylation levels regardless of treatment effect (compare teal lines between Figure 3A and B and between Figure S8A and B). Similarly, H3K4me3 (teal lines in Figure 3C and D) and chromatin accessibility (teal lines in Figure 3E and F) are also lower in genes upregulated following HDACi treatment, without substantial change in H3K4me3 due to treatment (compare between teal and pink lines in Figure 3C and D). However, promoter chromatin accessibility was particularly lower at downregulated genes due to treatment (compare between teal and pink lines in Figure 3E and F). Active nucleosome remodeling may contribute to these observations, where histones may be removed from promoters of upregulated genes and histones may be added to the promoters of downregulated genes (Figure 3G); therefore, this would present as reduced acetylation in upregulated genes given the nucleosome removal. We find that at 24 hpt approximately 23% of upregulated gene promoters possess differential OCRs with increased accessibility. Thus, if a promoter is otherwise accessible (nucleosome-free) to increase gene expression, nucleosome remodeling (histone replacement) may not be necessary. Conversely, approximately 62% of downregulated genes promoters possess differential OCRs with decreased accessibility suggesting that gene down regulation is more associated with nucleosome remodeling. Further, histone acetyltransferase genes were down regulated in response to HDACi treatment, which further corroborates our finding of reduced histone acetylation density at promoters, particularly at downregulated genes (Table S1). Nucleosome remodeling without active addition of histone acetylation may underlie the observed chromatin profile of differentially expressed genes.

**Figure 3:**
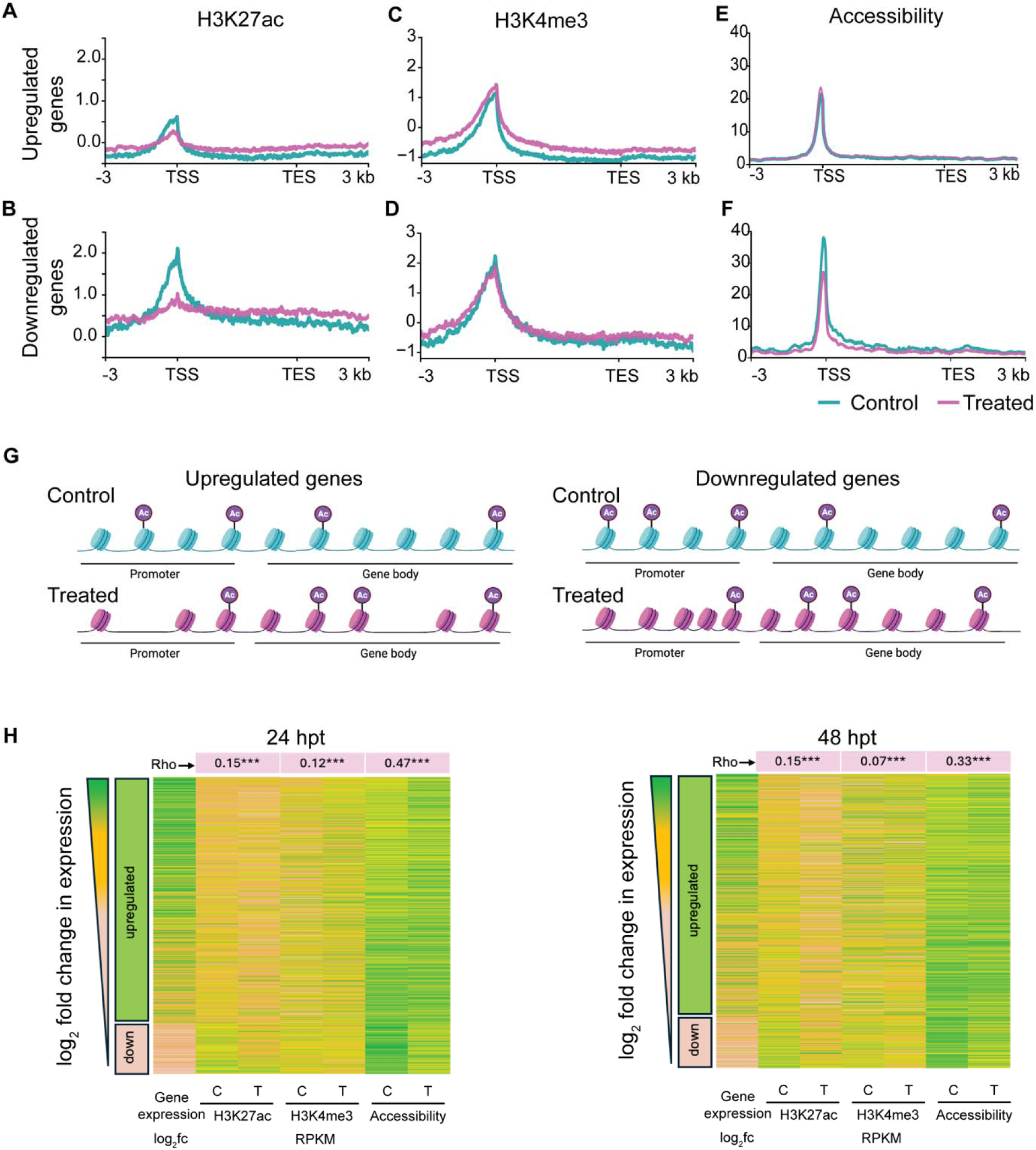
Nucleosome remodeling is associated with gene expression change. A,B) H3K27ac, C,D) H3K4me3, and E,F) accessibility profiles across upregulated and down regulated genes at 24 hpt with greater than 2-fold change. All genes are scaled to 5 kb and genes ± 3 kb regions are shown. Control (teal) and Treated (pink) profiles are overlaid. G) A speculative model presenting nucleosome remodeling at differentially expressed genes. H) Heatmaps representing log_2_ fold change in gene expression of significant differentially expressed genes at 24 hpt (left) and 48 hpt (right) (adjusted p-value <0.05 and log_2_ fold change>2) sorted in the order of positive to negative fold change in expression. RPKM values of promoter (TSS ± 500 bp) H3K27ac, H3K4me3, and accessibility in control and treated cells are shown in the subsequent columns for those genes. Spearman rank correlation (Rho) between the fold change in expression versus the fold change in each of the features is provided on top. *** indicates p-value <0.001.

At both timepoints, changes in gene expression due to treatment are moderately correlated with changes in promoter chromatin accessibility (Spearman’s rank correlation coefficient ρ = 0.47 at 24 hpt and ρ = 0.33 at 48 hpt). Gene expression changes have a weak positive correlation with changes in promoter H3K27 acetylation at 24 hpt (ρ = 0.15) and H3K4 trimethylation (ρ = 0.12) (Figure 3H). For those differentially expressed genes identified at 24 hpt, their expression at 48 hpt retains a weak significant positive correlation with the H3K27 acetylation profile at 48 hpt (ρ = 0.15), and a slight reduced but weak correlation with H3K4 trimethylation (ρ = 0.07) (Figure 3H right panel). Thus, changes in the nucleosome profile and epigenetic modifications directly following HDACi treatment are associated with changes in gene expression. Further, the retained gene expression changes at 48 hpt are still connected with promoter chromatin accessibility (ρ = 0.33), suggesting promoter nucleosome remodeling is largely maintained after HDACi treatment and associated with differential gene expression. Of note, changes in gene body features, except chromatin accessibility, are weakly negatively correlated with changes in gene expression (Figure S9). When changes in each of the features were compared between 24 and 48 hpt for promoters of significant differentially expressed genes, we find the highest positive correlation for gene expression (Spearmann’s ρ = 0.64) followed by chromatin accessibility (ρ = 0.61), then H3K27ac ((ρ = 0.27), and finally H3K4me3 (ρ = 0.17), All correlation coefficients are statistically significant with p-value <0.001. (Figure S10). As such, while early changes in gene expression and accessibility are largely retained even after 48 hpt, histone modifications (H3K27ac and H3K4me3) are relatively more dynamic despite the retention of functional change.

### Moderate bidirectional changes in 3D chromatin architecture are induced by HDACi treatment

Hi-C data provide information on the frequency and distribution of three-dimensional contacts among chromatin regions. While SAHA induced substantial changes in the epigenome profiles and gene expression, treatment did not impact higher order chromatin organization to the same extent at least at the time points we investigated. We found fewer short-range chromatin contacts (∼200 kb) and more long-range contacts (∼3 MB) in response to HDACi treatment at 24 hpt. This difference can be visualized in the lower black and white plot of Figure 4A, where the divergence from zero is plotted as Treated/Control demonstrating the effect of treatment. Similarly, differential chromatin contacts between control and treated cells at any distance were minimal at 48 hpt, thus contact frequency was not considerably different (Figure 4A, lower panel).

**Figure 4:**
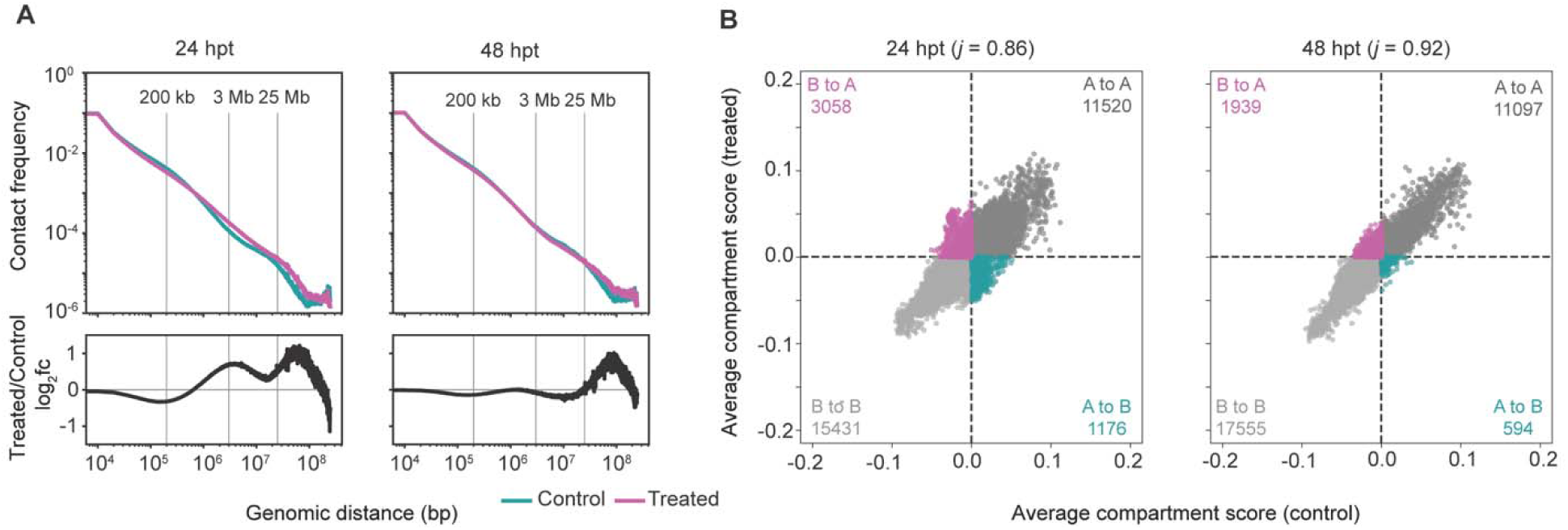
Moderate bidirectional change in 3D chromatin organization at 24 hpt subsides by 48 hpt. A) Contact frequency as a function of genomic distance in log10 scale is plotted for Control (teal) and Treated (pink) cells at 24 and 48 hpt. Replicates are combined. Bottom panel (black) shows the log_2_ fold change in contact frequency between Control and Treated samples as a function of genomic distance in log_10_ scale. B) Replicate averaged compartment scores of Control versus Treated cells at 100 kb windows across the genome are plotted as a scatter plot for 24 hpt and 48 hpt. Compartment exchange events are represented by off-diagonal points. Teal and pink represent A to B and B to A compartment exchange, respectively. Grey points represent compartments without a change in identity. The number of compartments in each category is annotated in the respective quadrant. The measure of overlap between Control and Treated compartment assignments is represented by the Jaccard index (j) given in the plot title.

At broader scales, chromatin is organized into active (A) and inactive (B) spatial compartments by segregating genomic elements for active transcription within A compartments (Harris et al., 2023). We defined chromatin compartments at 100 kb resolution using eigenvalues computed from Hi-C data and nucleosome accessibility information from ATAC-seq data (described in methods section: compartment analysis). Assessment of compartment exchange demonstrates the rearrangement of genomic structural features, which can vary (up to 60% exchange) among cell types and in response to perturbations (Schmitt et al., 2016). Thus, we compared compartment assignments between control and treated cells and found moderate genome-wide chromatin compartment exchange in response to HDACi treatment (Figure 4B). At 24 hpt, approximately 13.6% of the genome (100 kb regions) exchanged chromatin compartment identity with the majority (∼ 72%) switching from the inactive (B) to active (A) chromatin compartment, suggesting a moderate alteration of chromatin structure. At 48 hpt, approximately 8.1% of the genome (100 kb bins) switched compartment identity compared to Controls indicating a partial reversal of treatment-induced compartment exchange. For both genes with differential expression and OCRs with differential accessibility, ∼10% and 11% (respectively) overlap with an exchanged chromatin compartment (A to B or B to A). Thus, while HDACi induces rearrangement of genome structure, regulation of transcription and nucleosome remodeling is likely not entirely (nor linearly) dependent upon large-scale compartment exchange. Rather, several scale-dependent, locus-specific factors may influence the relationship between broad-scale chromatin compartment exchange and fine-scale functional changes.

### Chromosome and locus-specific association between compartment exchange and fine-scale functional features

To further investigate relationships among broad-scale structural features and fine-scale functional responses, we performed continuous wavelet transformation (CWT) of B-to-A compartment switching, gene upregulation, and gene density in response to HDACi treatment. To allow for a general comparison, we focused this analysis on a set of representative chromosomes (1, 17, 18, and 19) with particular characteristics, including variations in gene density. For reference, chromosome 1 is the largest (248.4 Mb) human chromosome with moderate gene density (22.6 genes/Mb). Chromosome 17 is small (84.3 Mb) and the second most gene-dense chromosome in the human genome (36.2 genes/Mb). Chromosome 18 is also small (80.5 Mb) but is quite gene-poor (15.5 genes/Mb), and chromosome 19 is even smaller (61.7 Mb) but is the most gene-rich chromosome in the human genome (46.9 genes/Mb). Aside from average differences in gene density across whole chromosomes, we also considered differences in the localization of gene-dense and gene-poor regions along these chromosomes (Figure S12).

Overall, analysis of CWTs for B-to-A compartment-switching and gene upregulation revealed chromosome-specific, location-specific, and scale-dependent signatures and correlations (Figure 5, Figure S11). These chromosome-specific patterns were found for both B-to-A compartment transition CWTs (Figure 5A and D, Figure S11 A and D) and gene upregulation CWTs (Figure 5B and E, Figure S11 B and E). Here, B-to-A compartment switching at broader scales, where the signal is aggregated along wider stretches of the chromosome, is comparable for chromosomes 1, 17, and 19 with higher signal density on the p-arm of the chromosome compared to the q-arm of the chromosome (Figure 5A and D, Figure S11 A and D). Conversely, on chromosome 18, the B-to-A compartment switching signal density spans the centromere and covers a large portion of the q-arm. As the wavelet becomes more contracted at finer scales (i.e., lower values), the signal is aggregated over smaller regions of the chromosome, and differences between chromosomes become more pronounced (Figure 5A and D, Figure S11 A and D). For gene upregulation, differences in CWT heatmaps across chromosomes are very evident even at broader scales (Figure 5B and E, Figure S11 B and E). These variations can only be explained in part by gene density patterns across chromosomes (Figure S12): gene upregulation and gene density CWT heatmaps show strong concordance in chromosomes 1 and 19, which are relatively gene dense. However, lower concordance between gene upregulation density and gene density was observed for both gene-dense chromosome 17 and gene-poor chromosome 18. Thus, chromosomal differences in B-to-A compartment switching and upregulation of gene expression are likely predicated on other chromosomal features in addition to gene density.

**Figure 5:**
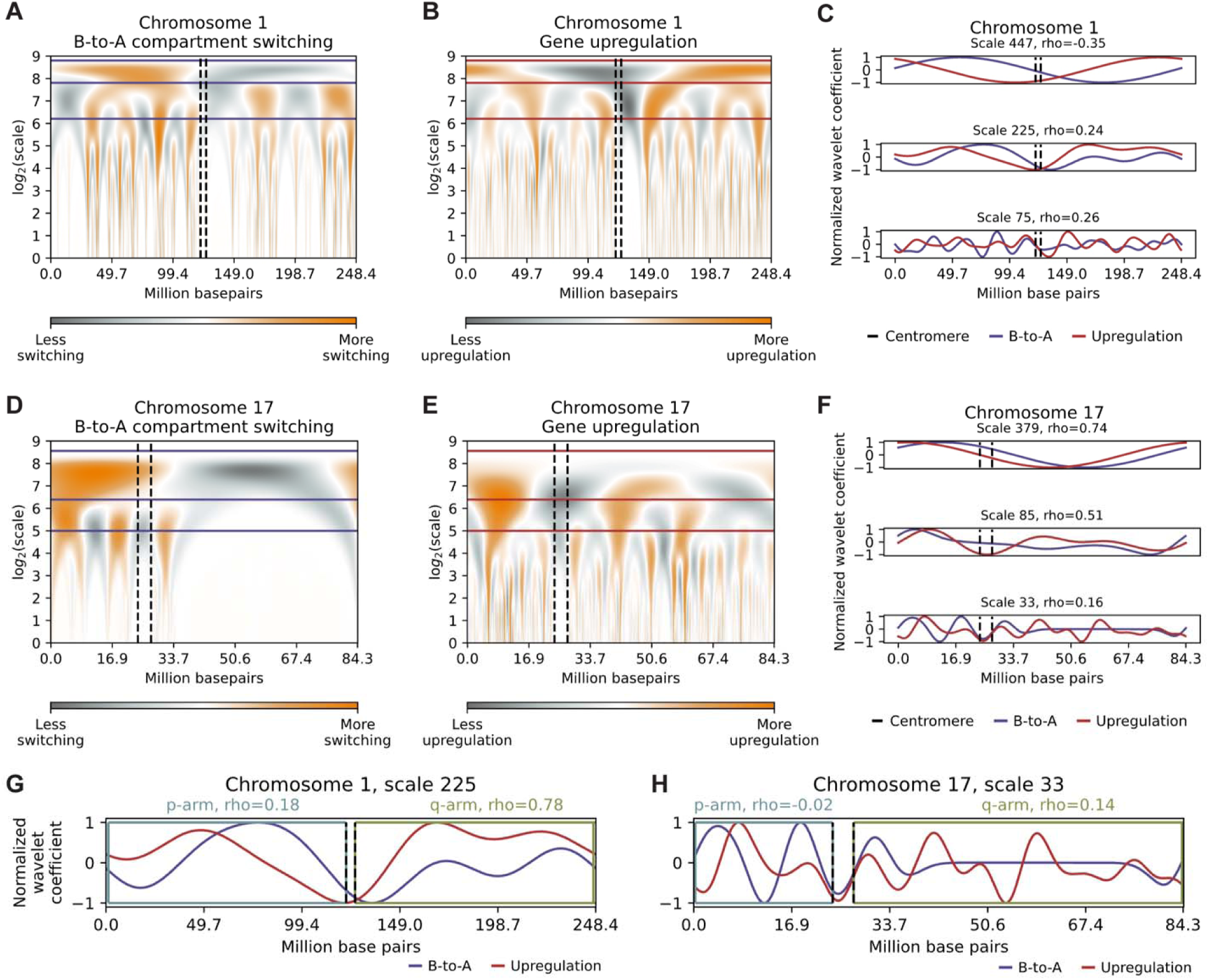
Concordance between B-to-A compartment switching and upregulation of gene expression are dependent upon chromosome and location within the chromosome. A) Density of B-to-A compartment switching signal at 100 kb windows along chromosome 1 using continuous wavelet transformation. B) Density of upregulated genes at 100 kb intervals along chromosome 1 using continuous wavelet transformation. C) The intersection of the B-to-A wavelet and the gene upregulation wavelet at different scales (447, 225, 7) on chromosome 1. The Pearson’s correlation coefficient (r) at each reported scale is provided in the title. D) Density of the B-to-A compartment switching signal at 100 kb windows along chromosome 17 using continuous wavelet transformation. E) Density of upregulated genes at 100 kb intervals along chromosome 17 using continuous wavelet transformation. F) The intersection of the B-to-A wavelet and the gene upregulation wavelet at different scales (379, 85, 33) on chromosome 17. The Pearson’s correlation coefficient (r) at each reported scale is provided in the title. G) Correlations between the B-to-A wavelet and the gene upregulation wavelet at scale 85 on the p arm (left) and q arm (right) on chromosome 1. H) Correlations between the B-to-A wavelet and the gene upregulation wavelet at scale 85 on the p arm (left) and q arm (right) on chromosome 17.

In addition to differences *within* structural features and functional responses across chromosomes, concordance *between* them also differs across chromosomes (Figure 5, Figure S11). Most strikingly, while particularly high concordance between B-to-A compartment switching (broad scale) and gene upregulation (function) is observed for gene-poor chromosome 18 (Figure S11A and B), no such concordance is observed for chromosomes 1, 17, and 19 (Figure 5A and B, Figure 5D and E, Figure S11D and E). Additionally, for all chromosomes, the levels of concordance between features depends upon the scale of the wavelet and may increase or decrease as scales become broader or finer (Figure 5C and F, Figure S11C and F), though the relative change and strength of correlation is highly chromosome dependent. Furthermore, in addition to scale dependency, concordance between B-to-A compartment switching and gene density may be predicated on chromosome regions as illustrated by the variations in correlation strength, which are determined by wavelet scale and localization on the p- or the q-arm of the analyzed chromosomes (Figure 5G and H, Figure S11G and H). Taken together, concordance *between* modalities is dependent upon chromosome region and the scale of analysis, where the strongest relationships vary across individual chromosomes.

### Genome-wide weakening of Topologically Associating Domains with pronounced contact reduction at downregulated genes

Next, we investigated the dynamics of fine-scale chromatin organizational features, including topologically associating domains (TADs) and chromatin loops (Mirny et al., 2019; Rao et al., 2014). Topologically associating domains (TADs) are genomic regions that spatially associate more than expected by chance (Dixon et al., 2012) (example shown in Figure 6A). When *cis* interactions within TADs (intra-TAD interactions) in control and treated cells at each time point were compared, we found an overall weakening of intra-TAD interactions at 24 hpt, demonstrated as negative values in (log_2_) fold change for contact frequency (Figure 6B). Interestingly, these weakened TADs regained their strength at 48 hpt (an increase in fold change), highlighting the dynamic nature of TAD organization (Figure 6B, Figure S13). The fold change among intra-TAD contacts among control cells was minimal when comparing across the time points (Figure 6B), while an increase in contact frequency was observed between 24 and 48 hpt in treated cells (Figure 6B). Thus, while intra-TAD contact frequency was reduced due to treatment, TAD boundaries are largely conserved at both timepoints (marginally different at 24 hpt) as we find similar insulation score profiles (Figure S14). Together, these observations suggest that SAHA treatment led to an initial weakening of intra-TAD interactions that were partially restored by 48 hpt without considerably affecting the TAD structure.

**Figure 6:**
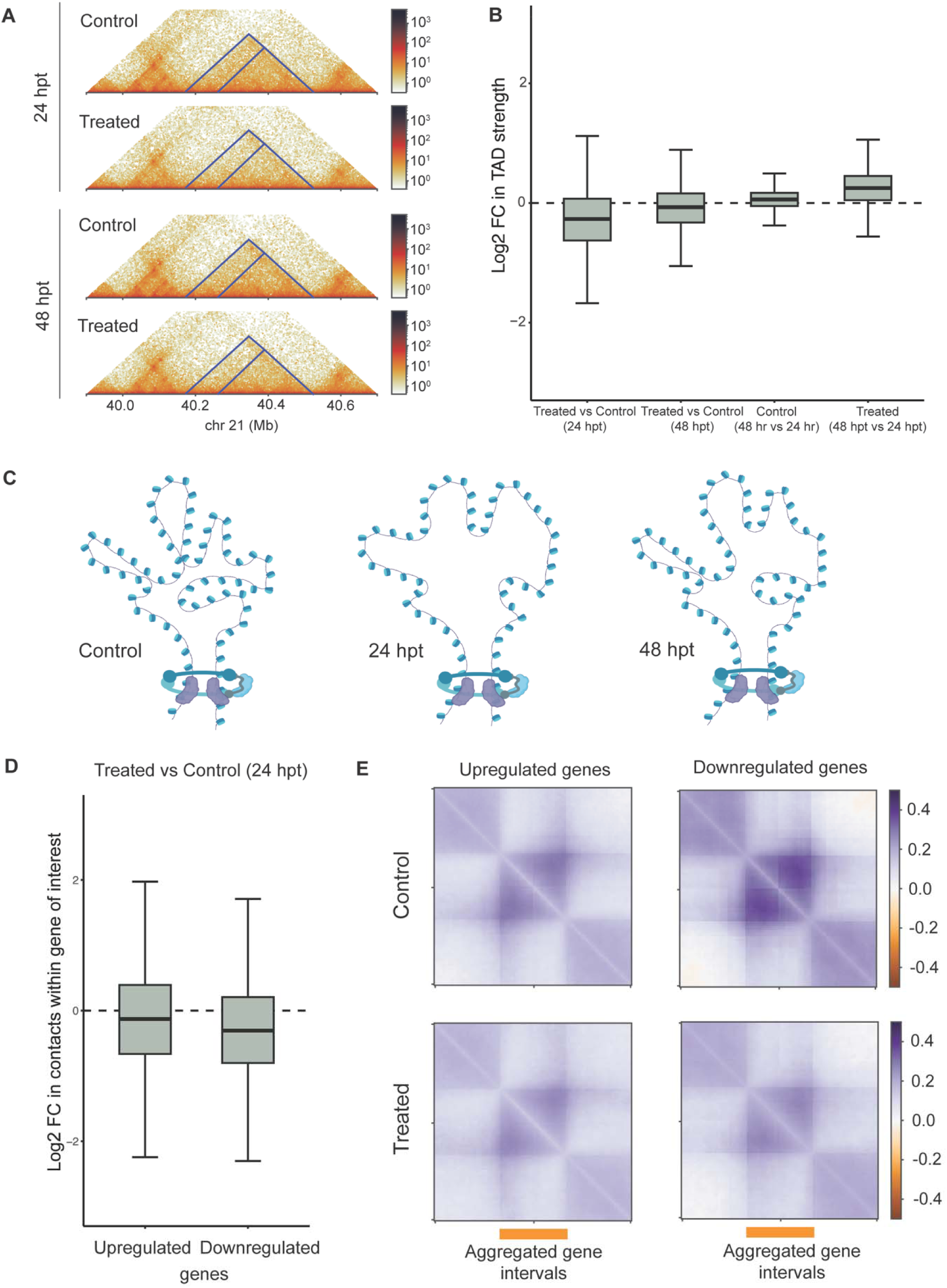
Moderate genome-wide weakening of Topologically Associating Domains with pronounced contact reduction at downregulated genes. A) Control and Treated Hi-C heatmap from 24 hpt and 48 hpt time points showing an example region comprised of dynamic TADs (triangular feature highlighted with blue lines) after HDACi treatment. B) Log_2_ fold change in observed/expected intra-TAD contact frequency are shown as box plots between i) treated and control cells at 24 hpt, ii) treated and control cells at 48 hpt, iii) control cells at 24 and 48 hpt, and iv) treated cells at 24 and 48 hpt. The central horizontal line denotes median TAD strength in each category. Outlier values are omitted from the plot. C) A schematic model representing possible intra-TAD weakening in treated samples at 24 hpt and partial regaining of contacts at 48 hpt. Loop boundaries are largely unchanged. D) Box plots summarize log_2_ foldchange in observed/expected contact frequency between treated and control cells at genes upregulated and downregulated at 24 hpt. The central horizontal line denotes median TAD strength in each category. Outlier values are omitted from the plot. E) Aggregated log_2_ fold change in observed/expected contact frequency (control: top panel, treated: bottom panel) across genes upregulated (left) and downregulated (right) at 24 hpt shown as heatmap. Contact strength within scaled, aggregated gene body is shown in the center (marked by the orange line underneath) with contact strength at matching sized flanking regions on either side. The heatmaps are symmetric representing same data on either side of the diagonal.

Further, to investigate the influence of intra-TAD contact reduction on gene expression, we assessed the change in contact frequency around differentially regulated genes and found a significant weakening irrespective of the direction of expression change (Figures 6D, E). The (log2) fold change in frequency for the majority of differentially expressed genes was below zero (Figure 6D). There are higher interactions at and around gene bodies suggesting that they are mostly found within TADs (shown as the observed/expected contact frequency heat map, Figure 6E). The shades of violet indicate higher than expected contact frequency (Figure 6E) in contrast to our observations of fewer than expected contacts outside TADs when TADs are aggregated genome wide (Figure S13, regions in shaded of orange). Curiously, the magnitude of contact weakening was stronger at downregulated genes (Figure 6D) as they possess higher contact frequency in control conditions (Figure 6E, top right panel). This suggests that either a subset of genes was more affected or was slower to recover from global TAD weakening that occurred after SAHA treatment.

Next, we identified chromatin loops as regions of increased contact frequency between loop anchor loci in a 2D Hi-C heatmap (Sanborn et al., 2015). While we detected an average of 14,000 fine-scale loops per replicate, and only a small number of them are statistically significant differential loops (Figure S15). This could be due to the heterogeneity across cells leading to weaker signals or due to the transient nature of functional chromatin loops. We combined replicates and identified several loops with considerable fold changes in interaction frequency (as exemplified in Figure S16A) suggesting dynamic loop reorganization. To further distinguish functional loops, we made a subset of loops called ‘promoter loops’, where at least one of the loop anchor intervals overlaps with a transcription start site. Promoter loops and contacts are thought to play a major role in regulating gene expression (Schoenfelder and Fraser, 2019; Zhang et al., 2022). We assessed the promoters of differentially expressed genes and found a weak, negligible correlation between the fold change in promoter contact frequency and the fold change in gene expression in a genome-wide level (Figure S16B). This correlation weakens when we assessed promoters that interact with potentially active enhancers (Figure S16C). Annotated enhancers for A549 cells (Gao and Qian, 2020) were deemed ‘active’ if they possess signals of accessibility including H3K27ac and H3K4me3 (Figure S17, described in methods section ‘Enhancer Annotation’). This suggests that at the genome-wide level the minor loop contact frequency change detectable at 24 hpt is not predictive of the differential gene expression.

After aligning all major chromatin features considered in this study (Figure 7), we find an example dynamic region with noticeable TAD boundary weakening at 24 hpt that partially regains the TAD insulation by 48 hpt. ((TAD marked with blue vertical dotted lines in Figure 7B, boundary weakening marked with arrow marks in Figure 7C.) Despite broadening and overall weakening of the histone acetylation profile across the entire region at both 24 and 48 hpt, the distribution of H3K4me3 is largely unaltered and still tightly regulated. However, two consecutive genes in this region (marked by the red-yellow arrows in expression track) increase in expression at 24 hpt and with a rebound (decrease) by 48 hpt. Concurrent with the gene expression changes, we found a slight reduction in accessibility and intra-TAD contacts more notably at 24 hpt (marked by blue dotted lines in Hi-C heatmap). While the directionality of relationship between chromatin structural changes (accessibility and contacts) with gene expression change can be context dependent, this example region presents concordance in gene expression and chromatin organization dynamics.

**Figure 7:**
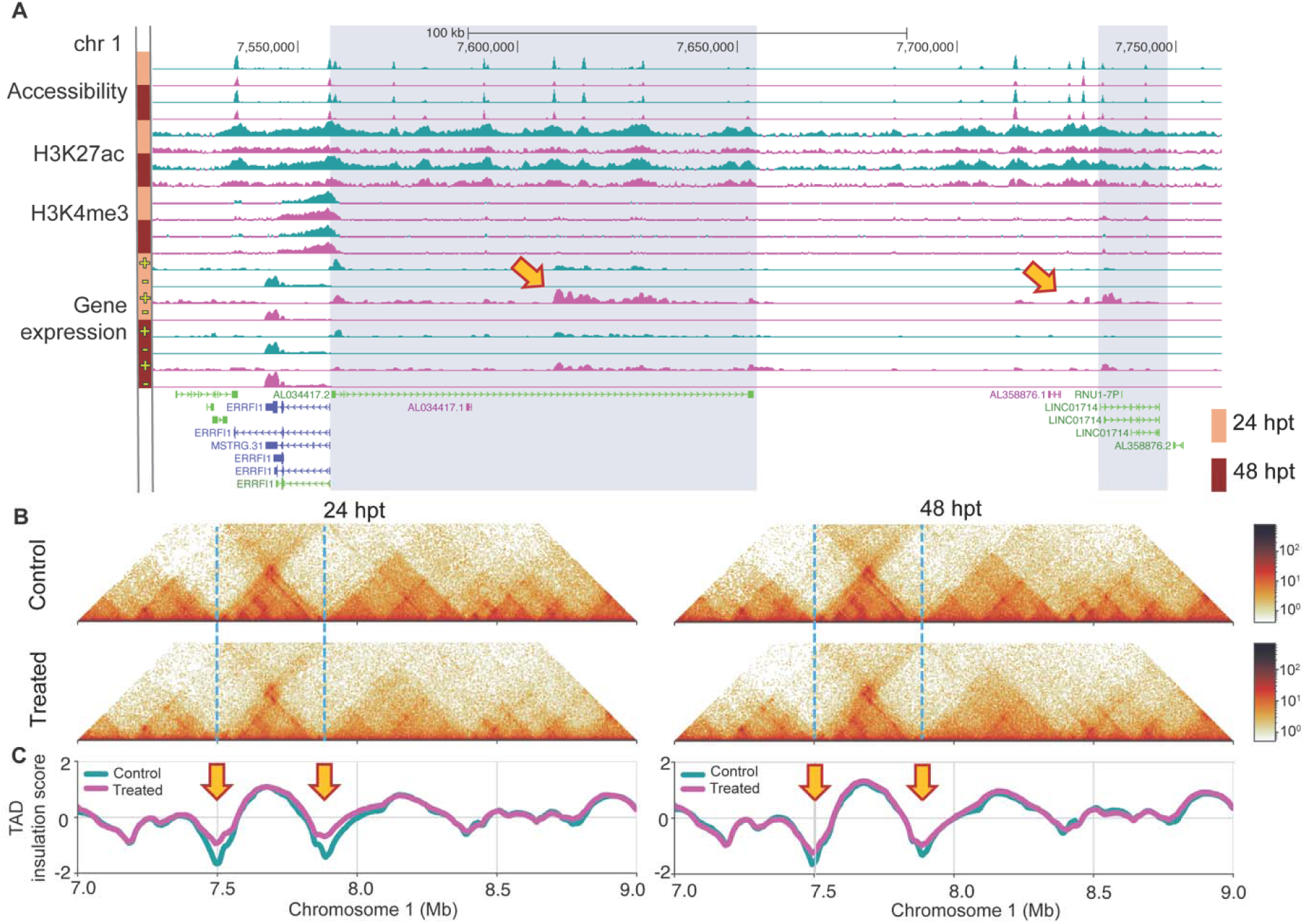
Concurrent alterations in TAD strength and gene expression. A) Replicate averaged profile tracks of chromatin accessibility, H3K27ac, H3K4me3, and strand specific gene expression are aligned for Control (teal) and Treated (pink) cells for a representative region in chromosome 1. Tracks for 24 hpt and 48 hpt are denoted with peach and maroon color bars respectively on the left-hand side. Annotated genes in this region are shown at the bottom most track. Red-yellow arrows point to the differentially expressed genes in this region. The region consisting of the differential genes are highlighted with gray shading. B) Hi-C heatmap of 24 hpt (left) and 48 hpt (right) control (top) and treated cells (bottom) of a broader region including the region of interest shown in panel A (highlighted with red rectangle). C) TAD insulation score for Control (teal) and Treated (pink) Hi-C maps shown in panel B. Arrows mark a TAD boundary with evident TAD weakening.

## Discussion

In this study, we present a comprehensive genome-wide investigation of chromatin structure-function changes following a genome-wide epigenomic perturbation. In general, HDAC inhibitors, and more specifically SAHA, elicit altered gene regulation resulting in curbed tumor growth, allowing cells to survive and adapt (Halsall et al., 2015). Given the extensive characterization of SAHA’s impact on gene expression, (Beckers et al., 2007; Halsall et al., 2015; Slaughter et al., 2021; Wu et al., 2015), we employed this system (SAHA in A549 cancer cells) to unravel the nature and persistence of locus-specific chromatin remodeling associated with genome function. Leveraging recent advancements in multi-omic experimental and computational techniques, we assessed how alterations in chromatin, particularly histone modifications, accessibility, and higher-order genome architecture, are associated with altered gene expression. Here, we demonstrate the anisotropy of genome architecture and chromatin features, which is predicated on time scales and unequal distribution of genome-wide impacts. Further, dynamic processes are not ubiquitous across all cells nor are patterns in features similar across all chromosomes, suggesting the need for cell- and chromosome-specific analysis. We find that locus-specific chromatin features, potentially including the rate of change in dynamics, are largely responsible for the direct influence of functional regulation. As such, targeted epigenetic therapies, either for specific loci or mechanisms that cause differential chromatin dynamics, could fine-tune these chromatin features to have minimal off-target effects with non-specific drug treatments.

In agreement with other studies that have employed SAHA treatment, we find elevated histone acetylation levels across the genome, particularly across gene bodies. This change in gene body acetylation suggests that HDAC enzymes play a key role in the maintenance of acetylation homeostasis. Indeed, previous work notes that increased H4 acetylation in gene bodies retargets bromodomain-containing acetylation readers, thereby affecting gene expression (Slaughter et al., 2021). However, we find increased gene body acetylation in both up and down-regulated genes in response to SAHA, suggesting that H3 acetylation in gene bodies does not directly correlate with gene expression changes. Additionally, there was reduced acetylation signal intensity at gene promoters after treatment. We also found several key lysine acetyltransferase enzyme genes were downregulated, corroborating previous reports of the same phenomenon (Halsall et al., 2015). Our novel observations suggest that SAHA not only impacts the ability to remove acetylation—via direct inhibition of histone deacetylase enzymes—but also impacts the mechanism for active, targeted histone acetylation via downregulation of histone acetyltransferase gene expression. This is likely a compensatory feedback mechanism to reduce the onslaught of aberrant hyperacetylation that occurs with HDAC inhibition. While we find the strongest changes are elicited 24 hours post treatment, there is a slight rebound effect as cells attempt to recover after this dramatic deviation in normality. Interestingly, acetylation levels do not rebound to the same extent as gene expression, further underscoring the non-linear relationship between the two. Finally, other chromatin features were impacted by SAHA to a lesser extent than acetylation, and therefore, may more directly support the moderate renormalization of genome functionality after treatment. Notably, the complete recovery to control levels of any chromatin feature or structure does not occur at least by 48 hours, suggesting some semi-permanent changes in response to treatment.

While the field generally finds that histone acetylation is associated with open chromatin regions (Bannister and Kouzarides, 2011), our understanding of the relationship between modifications and associated nucleosome occupancy is still evolving (Mansisidor and Risca, 2022). In this study, increased acetylation genome-wide did not constitute a matched increase in chromatin accessibility; rather, accessibility on a genome-wide level was mostly unscathed by SAHA treatment. Thus, perturbation in the acetylation equilibrium is not sufficient to completely override the balance of chromatin accessibility dynamics. Furthermore, of the small number of OCRs impacted by SAHA, there are nearly equal numbers with increasing or decreasing accessibility, though regions with prominent changes were more likely to be open than closed. Previous studies also report minimal bidirectional change in chromatin accessibility with increased (Baumann et al., 2021) or decreased (Hogg et al., 2021) global acetylation levels. While it seems that global acetylation changes do not directly correlate with global accessibility changes, the reduced expression of histone acetyltransferase enzymes resulting in reduced active acetylation of promoters may have affected the accessibility of several loci resulting in this observation. The majority of differentially accessible regions (71.5%) were associated with either genic regions or annotated enhancers highlighting that nucleosome remodeling largely occurred at functionally relevant genomic regions in response to treatment. These findings suggest that chromatin accessibility may be a pervasive feature that is more strongly connected to other features such as chromatin architecture and gene expression regulation than acetylation.

Global chromatin architecture delineated from Hi-C assessments was the least impacted by SAHA treatment compared to other chromatin features. While ∼13% of the genome exchanged compartments, only ∼10% of differentially expressed genes and ∼11% of OCRs were associated with compartment exchange. Thus, there is a disconnect between transcriptional regulation, nucleosome remodeling and large-scale compartments. While this assessment suggests weak associations at the global genome-wide level, our continuous wavelet transformation analysis demonstrates that concordance between chromatin architecture, gene upregulation, and gene density is specific to each chromosome, particular to genomic location, and dependent upon the scale of analysis. Indeed, differences in concordance were observed in different chromosomal arms and in chromosomes with more gene density; the concordance between modalities was also highly context dependent as well. Thus, discerning architectural changes at the genome-wide level may miss strong differential responses that occur on specific chromosomes, and even on specific regions of those chromosomes. To delineate the relationship between higher-ordered structure and genomic function, analysis on a per chromosome basis may be warranted. In addition to chromosome-specific responses, heterogeneity in cell culture may also underlie these context-dependent variations as well.

In addition, in assessing broader scale dynamics (compartment switching), we determined how fine-scale architectural features, including chromatin loops, respond to SAHA treatment. Loops are thought to facilitate interactions among gene promoters and enhancers or silencers, thereby directly impacting gene expression (Schoenfelder and Fraser, 2019; Zhang et al., 2022). However, the directionality of gene expression depends upon the gene’s position in the chromatin loop relative to the chromatin loop anchor (Reed et al., 2022). In our study, while SAHA induced minor changes in contact frequency at chromatin loop anchors, we found a weak positive correlation between differential gene expression and differential loops overlapping gene promoters. This observed weak correlation between differential enhancer-promoter loop contacts and differential gene expression may result from the faster turnover of three-dimensional chromatin contacts relative to gene expression. Potentially, we may have not captured the entirety of chromatin reorganization associated with functional changes at 24 hours post treatment, particularly given that the number of differential loops is even further reduced at 48 hours post treatment. Additionally, promoter interactions, including contacts with other regulatory elements such as silencers, may play a regulatory role in gene expression (Zhang, See, Tergaonkar, & Fullwood, 2022). Loop anchors may associate with TAD boundaries, which denote regions of highly interactive chromatin, based on the number and intensity of their chromatin contacts determined from Hi-C analysis. In conjunction with our differential loop analysis, we found weakening of TAD interactions, suggesting loss of structural integrity or insulation between adjacent regions. We and others (Venu et al., 2023; Wang et al., 2023) have observed this phenomenon in response to other stressors, including viral infection. However, in this study, TAD weakening occurred around both up- and down-regulated genes but was slowly regaining by 48 hours post treatment. In the example region presented (Figure 7), the weakening of TAD strength at 24 hours post treatment was associated with increased expression of more than one gene in the region. Further, expression of those genes decreased (closer to control levels) with regaining of TAD strength at 48 hpt. While the directionality cannot be generalized from our data, this observation indicates an association between gene expression changes and chromatin architectural turnover, where turnover may occur more rapidly than expression change.

Among the chromatin features analyzed in our study, we found the most compelling and discernable differences occurred at downregulated genes after SAHA treatment. The promoters of downregulated genes had significant decreases in histone acetylation *density* and chromatin accessibility, suggesting nucleosome remodeling included the addition of new histones to promoters. While global acetylation increased, new histones were not acetylated, likely due to reduced histone acetyltransferase expression, and thus acetylation density was reduced (rather than a pure reduction in acetylation). Conversely, in promoters of upregulated genes, there was a marginal increase in chromatin accessibility with largely unaltered histone acetylation density profiles, suggesting nucleosome remodeling removed histones. It is uncertain whether these phenomena are the result of transcription or an active process involving chromatin remodelers. Further, in genes upregulated due to SAHA treatment, the lower basal levels of acetylation and accessibility at their promoters suggests that those regions are highly dynamic. In this case, this lower level is likely an artifact of the experimental assay, where the strength of the signal is determined by the stability of the epigenetic modification in thousands to millions of cells. That is, when dynamic chromatin remodeling occurs rapidly, not all cells will present the same signals at the same time (Lai and Pugh, 2017). Thus, the investigation of dynamic (versus stable) features will have representation from fewer cells and therefore a weaker overall signal than a stable feature from the same collection of cells. In line with this, smaller differences in TAD strength due to treatment for upregulated genes (compared to downregulated genes) also indicates that these regions are undergoing dynamic chromatin restructuring. The observed global TAD weakening could be a genome-wide response to global HDACi inhibition; however, regions with faster turnover of chromatin remodeling mechanisms adapted allowing for permissive gene transcription. Conversely, genomic regions with slower chromatin turnover possibly present a less permissive state. These findings are particularly of interest as perturbation in steady-state histone acetylation equilibria in either direction have shown similar effects on gene transcription, chromatin accessibility and 3D chromatin architecture (Hogg et al., 2021).

Overall, our study presents an in-depth investigation of multiple chromatin features after major epigenome perturbation with a clinically relevant cancer drug. While it is reassuring to find the resilience of the structural and functional landscape of the genome despite global alterations in the histone acetylation profile, our study unravels the correlation between locus-specific chromatin dynamics and gene expression and finds some effects persist. We find stronger signals of non-permissive chromatin conditions at downregulated genes potentially arising from larger cell subpopulation suggesting that those regions are less dynamic compared to regions with upregulated genes. This finding necessitates future investigation of mechanisms underlying locus-specific rates of chromatin remodeling at the nucleosome level and at the level of higher-order organization at sub-megabase resolution. Given that stable chromatin features are more likely to occur in larger cell populations than dynamic features, single cell characterization of cellular heterogeneity would enable categorization of genomic regions based on their chromatin dynamics. Further, live cell imaging techniques are required for obtaining direct evidence of locus-specific chromatin dynamics. A clear understanding of regions that are potentially highly affected because of their slow chromatin dynamics and underlying mechanisms will benefit the design of therapeutics with minimized off-target effects.

## Methods

### Cell Culture

A549 cells, isolated from the lung tissue of a 58-year-old Caucasian male with lung cancer, were obtained from ATCC (Cat. #CCL-185). Cells were maintained in Complete Medium consisting of Dulbecco’s Modified Eagle’s Medium (DMEM) supplemented with 4 mM L-Glutamine (ATCC, Cat. #30-2002) and 10% (v/v) Fetal Bovine Serum (FBS, ATCC, Cat. #30-2020). Cells were incubated at 37°C with 5% CO_2_ and 95% relative humidity. Multiple vials were cryopreserved at passage #2 in liquid nitrogen vapor until needed for cell culture experiments. For cryopreservation, cells were resuspended to a density of 10^6^ cells per mL in ice-cold Freezing Medium consisting of DMEM, 20% (v/v) FBS, and 5% (v/v) dimethylsulfoxide (DMSO, ATCC, Cat. # 4-X).

For cell culture experiments, A549 cells were propagated for at least two additional passages prior to seeding for HDACi drug treatments (described below). Cells were washed once in 1X PBS (Gibco, Cat. # 10010049), detached using 1X TrypLE Express Enzyme (Thermo Scientific, Cat. # 12604021), resuspended into Complete Medium, and counted using the Scepter 3.0 Handheld Cell Counter and 60 µm Counter Sensors (Millipore, Cat. #PHCC360KIT). Cells were seeded for HDACi drug treatments at passage #7-8 into replicate tissue culture flasks (surface area = 175 cm^2^) in three biological replicates using 5−10^6^ cells per flask. Seeded flasks were incubated for 72 hrs at 37°C, yielding a confluent monolayer (∼10−10^6^ cells per flask) at the time of drug treatment.

### HDACi Drug Treatments and Cell Harvest

Cell-seeding medium was removed after 72 hrs from replicate tissue culture flasks and replaced with Complete Medium containing 10 µM of the HDACi drug suberoylanilide hydroxamic acid (SAHA) or containing an equivalent volume of the drug vehicle only, DMSO (Pierce, Cat. #85190). Mock-treated (control) and drug-treated culture flasks were incubated in parallel for 24 or 48 hrs at 37°C. During cell harvest (at 24 and 48 hrs post-treatment), drug-treated and mock-treated culture flasks for each biological replicate were processed in parallel for Hi-C, ChIP-seq, and RNA-seq analyses (described below). Cells were washed once in 1X PBS, detached using 1X TrypLE Express Enzyme for 20 min, then resuspended in Complete Medium to create a uniform single-cell suspension. Cell suspensions were held at room temperature and counted using the Scepter 3.0 Handheld Cell Counter and 60 µm Counter Sensors, then immediately divided as needed for downstream processing.

### Chromatin Conformation & Epigenomics Assays and Library Preparation

#### Hi-C

For Hi-C analysis, 1−10^6^ cells per aliquot were processed in triplicate 24 and 48 hrs after treatment as previously described (Venu et al., 2023). Briefly, cells were crosslinked with 37% (w/v) formaldehyde containing 10-15% (v/v) methanol (Sigma, Cat. #252549), treated with 200 mM glycine (Sigma, Cat. #G8790) and snap frozen for storage. As previously described, frozen crosslinked cells were processed into Hi-C libraries per the Arima Hi-C Kit manual (Cat. #A160134), the Arima-Hi-C Library Preparation user guide (Cat. # A160141), and the Accel-NGS® 2S Plus DNA Library Kit (IDT, Cat. #10009877). Libraries were sequenced on an Illumina NextSeq 2000 (PE 2−150bp) at the LANL Genome Center.

#### ChIP-seq

For ChIP-seq, 10−10^6^ cells per aliquot were processed in duplicate 24 and 48 hrs after treatment. Counted cells were resuspended in 1X PBS and crosslinked for 5 min using Pierce^TM^ 16% (w/v) methanol-free formaldehyde (ThermoScientific, Cat. #28906) at a final concentration of 1% (v/v). Formaldehyde was quenched with 125 mM glycine. Crosslinked cells were pelleted and washed twice in ice-cold 1X PBS (Gibco, Cat. #10010049). Chromatin was isolated and fragmented using the Diagenode Chromatin EasyShear Kit Ultra Low SDS (Cat. #C01020010) following the manufacturer’s protocol and Diagenode Bioruptor Pico (Cat. #B01080010) (30 sec on and 30 sec off settings for 17 cycles) to generate ∼350 bp sized fragments. Fragmented chromatin was stored at −80°C until use. Frozen chromatin was thawed on ice, and the volume was brought to 500 µl by adding IP dilution buffer (16.7 mM Tris-HCl, 0.01% SDS, 1.1% Triton X 100, 1.2 mM EDTA, 167 mM NaCl) supplemented with 1X Halt™ Protease Inhibitor Cocktail (Thermo Scientific, Cat. #78430). Fragmented chromatin was precleared with 15 µl Pierce Protein A/G beads (Thermo Scientific, Cat. #88803) at 4°C for 1 hr with rotation; 50 µl of the precleared chromatin was saved as input and stored at −80°C until use. The following antibodies were used for ChIP experiments: H3 (Abcam, Cat. #ab1791), H3ac (pan-acetyl) (Active Motif, Cat. #39139), H3K4me3 (Active Motif, Cat. #39060), and H3K27ac (Abcam, Cat. #ab4729). ChIP antibodies (4-5 µg) were prebound to 30 µl Protein A/G beads in 500 µl RIPA buffer (10mM Tris-HCl, 0.1% SDS, 1% Triton X 100, 1mM EDTA, 140mM NaCl, 0.1% Na deoxycholate) incubated for 2-4 hrs at 4°C with rotation. Conjugated beads were pulled down and washed 3X with RIPA buffer. Antibody-conjugated beads were added to precleared fragmented chromatin and incubated o/n at 4°C with rotation. Antibody-bound precleared fragmented chromatin was incubated with 30 µl Protein A/G beads for 2 hrs at 4°C with rotation. Complexes were pulled down, unbound lysate was removed, and beads were washed as follows for 10 min each at 4°C with rotation: RIPA; 2X RIPA 500 (RIPA buffer with 500 mM NaCl); LiCl buffer (10 mM Tris-HCl, 250 mM LiCl, 0.1% SDS, 1% NP40,1 mM EDTA, 0.5% Na deoxycholate); and TE buffer (10mM Tris pH 8, 1 mM EDTA). Chromatin was incubated in elution buffer (1% SDS, 0.1 M NaHCO_3_) at 65°C for 10 mins, incubated at RT for 10 min, and incubated o/n at 65°C. The input DNA was thawed and its volume brought to 300 µL with elution buffer. Both chromatin and input samples were treated with 40 µg/ml RNaseA (Qiagen, Cat. #19101) at 37°C for 30 min, followed by protein digestion with 80 µg/ml Proteinase K (ThermoScientific, Cat. #25530049). Samples were incubated with NaCl (100 mM final concentration) for 1 hr at 65°C in an Eppendorf thermomixer at 300 rpm. The DNA was purified using the Qiagen PCR cleanup kit (Qiagen, Cat. #28104), quantified via Qubit, and concentrated with Ampure beads (Beckman, Cat. #A63881) to remove any PCR inhibitors and meet library prep volume requirements. Libraries were generated via the Tecan Ovation V2 Ultra Low Kit with unique dual indexes (Tecan, Cat. #9149-A01) and sequenced on an Illumina HiSeq 2500 (PE 2−150bp) at the LANL Genome Center.

#### RNA-seq

For RNA-seq, 2−10^5^ cells were cryopreserved in Freezing Medium into duplicate 1 mL aliquots 24 and 48 hrs after treatment. RNA was extracted using the RNeasy UCP Micro Kit protocol kit (Qiagen, Cat. #73934) as previously described (Venu et al., 2023). Aliquots were thawed in treatment pairs (mock-treated and SAHA-treated) in a 37°C water-bath, transferred to ice-cold Complete Medium, then centrifuged at 1000 rcf for 2 min. Cells were washed in 10 mL ice-cold 1X HBSS (Gibco, Cat. #14025092) and lysed in 350 µL ice-cold RULT Buffer containing 1% (v/v) b-mercaptoethanol (Sigma, Cat. #63689). RNA extractions, performed immediately after cell lysis, and included an on-column DNase I treatment step for 15 min to deplete DNA. RNA was eluted in 20 µL RNase-free water and total RNA was quantified with the Qubit RNA HS Assay Kit (Thermo Scientific, Cat. #Q32855). RNA quality was measured using the Bioanalyzer RNA 6000 Assay (Agilent, Cat. #5067-1511). Ribosomal RNA was depleted from 500 ng total RNA using the Ribo-Zero Plus rRNA Depletion Kit (Illumina, Cat. #20037135). Sequencing libraries were generated using the Illumina NextSeq 500 High Output Kit v2.5 (Cat. #20024908) and eluted in DNA Elution Buffer (Zymo Research, Cat. #D3004-4-10). Library concentration and average size was verified using the TapeStation DNA 5000 Assay Reagents and Screen Tapes (Agilent, Cat. #5067-5589 and #5067-5588). Libraries were normalized and pooled based on the TapeStation concentrations, then pooled libraries were quantified using the Illumina/Universal Library Quantification Kit (KAPA Biosystems, Cat. #KK4824). Libraries were sequenced on an entire lane of NextSeq 500 HO flow cell to generate paired-end 151 bp reads (300 cycles).

#### ATAC-seq

In a separate experiment, A549 cells were cultured, seeded into flasks, and treated with 10 µM SAHA, and harvested exactly as described above. Approximately 2−10^5^ cells were cryopreserved in triplicate 1 mL aliquots using ice-cold Freezing Medium 24 and 48 hrs after treatment as previously described (Roth et al., 2023). Aliquots were subsequently thawed into Complete Medium, counted to achieve ∼80,000 cells per sample, and processed for ATAC-seq analysis using the Active Motif ATAC-seq Kit manual (Cat. #53150) as previously described (Roth et al., 2023). Briefly, cells were lysed and tagmented DNA was prepared and purified; DNA libraries were prepared via PCR using primers provided in the kit. Libraries were purified and additional size-selected using AmpureXP beads (Beckman, Cat. #A63880) per the manufacturer’s protocol to retain fragments between 150 bp and 600 bp. Libraries were quantified using a high-sensitivity Qubit dsDNA Quantification Assay Kit (Thermo Scientific, Cat. #Q32851) and a KAPA Library Quantification Kit (Roche, Cat. #KR0405). Libraries were sequenced via the Illumina NextSeq 2000 sequencer (LANL Genome Center) in paired-end mode (PE151) using P3 chemistry.

### Histone Extraction and Quantification

Cell lysis and subsequent histone extractions were performed using the Abcam commercial Histone Extraction Kit (Abcam, Cat. #ab113476) per the manufacturer’s instructions. At 24 and 48 hours post treatment, 5−10^6^ cells per culture (SAHA-treated and mock-treated) were lysed in 500 µL ice-cold 1X ‘Pre-Lysis Buffer’ and immediately snap-frozen in liquid nitrogen for long-term storage. Total histone proteins were purified from whole cell lysates and quantified using a Qubit Protein Assay Kit (Invitrogen, Cat. #Q33211) and the Qubit 2.0 Fluorometer. The Fluorometer was calibrated prior to each assay using the Qubit Protein Standards (provided in kit). Qubit Protein Assay results were corroborated using Bradford 1X Dye Reagent (BioRad, Cat. #5000205) and a dilution series Calf Thymus Histone H3 Standard (Roche, Cat. #11034758001) in distilled water. Samples and standards were incubated with 1X Dye (1:1) for 5 min, then absorbance was read immediately using a spectrophotometer at 595 nm. A commercial H3K27 Acetylation ELISA Kit (Active Motif, Cat. #53116) was used to quantify histone H3 acetylation induced by SAHA treatment on the extracted histones. Total input protein for each ELISA was normalized across all histone extracts to achieve 2 µg histone protein per well. Histone H3 protein standard (provided in kit) was used in a dilution series was used per plate; all conditions were assayed in triplicate. Absorbance was read using a spectrophotometer at 450 nm within 5 min of adding the Stop Solution.

### Microscopy

Approximately one million cells from each A549 culture harvested 24 and 48 hours post treatment were seeded into 6-well tissue culture plates with Complete Medium. Cells were incubated at 37°C with 5% CO_2_ and 95% relative humidity for 24 hrs. Cells were washed once in 1X HBSS, crosslinked with 4% (w/v) formaldehyde solution in PBS (Invitrogen, Cat. #FB002) for 15 min, and then washed twice with 1X HBSS. For nuclear staining, cells were incubated with DAPI Solution (Invitrogen, Cat. #62248) at 1µg/ml in PBS for 10 min and washed once in 1X HBSS. Cells were imaged using an EVOS M5000 Imaging system (Thermo Scientific, Cat. #AMF5000).

### Transcriptomics Analysis

We generated 12 RNA-seq libraries and sequenced to approximately 17 million read pairs per library. Raw read quality control was performed using FastQC (Andrews, 2010) and SeqMonk (Andrews, 2007) to verify library quality and degree of PCR duplication; all metrics are within the expected range. The human T2T reference genome and annotations were retrieved from NCBI. Annotations were converted to gtf format using the AGAT (Jacques Dainat, 2023) package function (agat_convert_sp_gff2gtf.pl). A splice junction aware reference index was generated for the combined genome and annotation files using STAR (Dobin et al., 2013) with the arguments (--runMode genomeGenerate --sjdbOverhang 149). Pre-alignment read trimming was not performed in favor of soft-clipping/quality filtering by the STAR aligner. Reads were aligned to the combined genome using STAR. Duplicate reads were marked but retained in downstream analysis per DESeq2 guidelines. Sense strand reads aligning to gene features were counted using htseq-count (Putri et al., 2022) with the arguments (--stranded=reverse). Antisense strand reads aligning to gene features were counted using htseq-count with the arguments (--stranded=yes). Differential expression analysis was performed using the DESeq2 (Love et al., 2014) matrix design (∼time + treatment). For visualization, replicate alignment files were merged using samtools. bigWigs were then generated using the bamCoverage function from the deepTools (Ramirez et al., 2016) package with the arguments (--effectiveGenomeSize 3117292070 --normalizeUsing BPM –filterRNAstrand {forward/reverse/NULL} --binSize 1).

### ATAC-seq Analysis

ATAC-seq libraries were sequenced as described above and analysis was performed as previously described (Roth et al., 2023; Venu et al., 2023). Twelve ATAC-seq libraries were sequenced to a median depth of 57 million read pairs per library. Briefly, raw reads were trimmed and filtered to remove Nextra adaptors and reads with repetitive sequences using Fastp (Chen et al., 2018). Bases with low quality scores (q < 15) were removed. Processed reads were aligned to the telomere-to-telomere (T2T) human reference genome, version 2 (Nurk et al., 2022) via BWA (Li and Durbin, 2009). Duplicate sequenced pairs and mitochondrial reads were marked and removed from analysis via Samblaster (Faust and Hall, 2014). Samples were further filtered using samtools with the following flags: -F 4 -F 256 -F 512 -F 1024 -F 2048 -q 30. Samples were then passed to MACS2 in BAMPE mode. Loci displaying significant enrichment of paired-end reads were identified as peaks using MACS2 (Gaspar, 2018; Zhang et al., 2008). Replicate correlation was assessed using deepTools (Ramirez et al., 2016) -plotCorrelation function on SPMR normalized MACS2 output big wig files. Open chromatin regions (OCRs) were defined by MACS2-called peaks followed the BEDtools (Quinlan and Hall, 2010) merge function. OCR differential accessibility analysis was performed using the DESeq2 (Love et al., 2014) matrix design (∼time + treatment).

### ChIP-seq Analysis

Raw read processing, mapping, and read filtration post mapping were performed for all ChIP and input reads as described for ATAC-seq data. Each ChIP-seq library was sequenced to approximately 23 million read pairs. For all ChIP experiments, high quality reads were analyzed using MACS2 to identify peaks with read enrichment. In contrast to ATAC-seq peak calling, corresponding treatment and time matched input control was used with each of the ChIP data.

### Chromatin Contact Map Generation

Hi-C libraries were sequenced as described above and analysis was performed as previously described (Venu et al., 2023). Eight Hi-C libraries were sequenced to have at least 600 million (150 bp) paired reads per library. Briefly, paired Hi-C reads were analyzed via Juicer Hi-C pipeline (Durand et al., 2016) and aligned to the T2T reference genome (Nurk et al., 2022). Restriction sites were identified, and Hi-C (.hic) files were generated using Juicer default settings with a mapping quality threshold of ‘q = 30’. We detected at least 700 million valid Hi-C contacts per condition. Replicate Hi-C maps were merged via Juicer (“mega” function).

### Replicate Correlation Analysis

For ChIP-seq, ATAC-seq, and RNA-seq datasets, replicate correlation was assessed using deepTools (Ramirez et al., 2016) -plotCorrelation function. SPMR normalized MACS2 output bigwig files were used as input for ChIP-seq and ATAC-seq datasets. For RNA-seq datasets, BPM normalized bigwig files generated by deepTools bamCompare function (parameters described above) were used. For Hi-C datasets, chromosome wise estimation of spectral statistic (Yan et al., 2017) was utilized to measure reproducibility between Hi-C replicates. For each data set, all the following analyses were performed on separated replicates unless otherwise mentioned; however, given high replicate concordance (Figures S2-S5), biological replicates were merged for visualization.

### Principal Component Analysis

For ChIP-seq, ATAC-seq, and RNA-seq datasets, principal component analysis was performed using the variance stabilization transformation package followed by plotPCA function from DESeq2. Raw reads from all individual samples across consensus peaks (i.e., merged intervals of MACS called peaks) per feature was used an input for ChIP-seq and ATAC-seq datasets. For RNA-seq datasets, read count per gene across all samples was used as input. The PCA for Hi-C datasets was conducted in Python using the FAN-C package at 50 kb resolution using --strategy fold change.

### Differential Peak Calling & Gene Expression

For ATAC-seq, H3K27ac, and H3K4me3 ChIP-seq datasets, differential peaks were called using DEseq2 with ∼time + treatment matrix design. Similarly, differentially expressed genes were also identified using DESeq2.

### Global and Local Profiling of Histone Modifications and Chromatin Accessibility

To determine the global levels of feature profile distribution in control vs treated samples, the deepTools –computeMatrix function with –binsize 10 kb was used. Input-normalized read coverage around midpoints of corresponding consensus peaks (i.e., merged intervals of MACS called peaks) were calculated for H3K27ac and H3K4me3 ChIP datasets. For ATAC-seq datasets, SPMR-normalized read coverage around OCR midpoints were calculated. For visualization, plotHeatmap and plotProfile functions from deepTools with or without kmeans clustering were used. Treated and control curve comparison plots were made and Wilcoxon rank-sum significance test was performed using a custom python script. FDR 2-stage Benjamini-Kriger-Yekutieli adjusted p-values are plotted. For plotting H3K27ac and H3K4me3 profiles normalized to histone H3 levels, input normalized H3K27ac and H3K4me3 profiles were further normalized to corresponding input normalized H3 data. Samtools merge and index functions were used to merge replicate bam files. deepTools bamcompare function was used for input normalization, and the bigwig compare function was used for H3 normalization. The feature profiles at promoters or gene bodies of differential expressed genes (padj <0.05 and absolute fc >2) were computed. The promoter was defined as TSS ± 500 bp region and the gene body was defined as the region between TSS and TES, excluding the promoter region. All regions of interest were scaled to 1 kb and the total H3 normalized (for ChIP data) and SPMR normalized (for ATAC-seq data) read count within each region was calculated.

### Distance Decay Analysis of Chromatin Contacts

Decay frequencies in contacts across genomic distances was performed using the FAN-C package (Kruse et al., 2020) in Python. Decay profiles for each chromosome at 10 kb resolution were calculated and combined for visualization and analysis.

### Loop Identification

HiCCUPs (Durand et al., 2016) within Juicer was used to identify chromatin loops at 5, 10, and 25 kb resolution across replicated and merged Hi-C maps. Hi-C contact counts were gathered for each replicate across the merged list of loops. From these counts, differential loops were determined via DESeq2 (Love et al., 2014), where comparisons among treatment (mock-treated and HDACi treatment) and time (24 and 48 hrs) were performed with time-matched controls per condition.

### Compartment Analysis

Compartment assignment scores at 100 kb resolution were estimated via Juicer (eigenvalue function). These scores combined with ATAC-seq profiles were used to label active (A) and inactive (B) compartments using the convention that the active A compartment has more ATAC-seq peaks. Compartment scores were reoriented (i.e., multiplying scores per chromosome by negative one) if the 95^th^ percentile of counts of ATAC-seq peaks were labeled as compartment B in a majority of ATAC-seq libraries. To identify compartment exchange due to SAHA treatment, we selected 250 kb bins assigned to the same compartments in both Hi-C biological replicates. The replicate average compartment score was determined for each bin only considering bins in agreement between replicates. The changes from positive (mock-treated) to negative (HDACi treated) bin score value was assigned as A to B exchange; similarly, changes from negative (mock-treated) to positive (HDACi treated) bin score value were assigned as B to A exchange. Bins that remained positive or negative were assigned as ‘no exchange’ A or B compartments, respectively.

### Enhancer Annotation

Cell type (A549) specific enhancer annotation was obtained from EnhancerAtlas 2.0 (http://www.enhanceratlas.org/). UCSC LiftOver tool was then used to convert enhancer coordinates in hg19 to T2T assembly. Potentially active enhancers were annotated by employing deepTools computeMatrix to compute control and treated chromatin accessibility, H3K27ac, and H3K4me3 signal across all scaled enhancers ± 1kb region. Further, enhancer clustering was performed on the resultant matrix with -kmeans 5. The enhancers that clustered together due to signals of higher accessibility, H3K27ac, and H3K4me3 in either control or treated samples were annotated as putatively active enhancers in the samples under study. A total of 10525 out of 49781 annotated enhancers were deemed active.

### Data preparation for continuous wavelet transformation

Continuous wavelet transformation was performed for gene density, B-to-A compartment switching, and gene upregulation in response to HDACi treatment. The B-to-A compartment switching signal is a binary indicator of whether B-to-A switching (a change from closed to open chromatin compartments) occurred in any given non-overlapping 100kb genomic bin. Gene density within each 100kb bin was computed by counting the number of genes, consisting of pseudogenes, non-coding RNAs, and protein-coding genes as reported in the T2T reference genome. The mean log_2_ fold change values of all significantly upregulated genes (FDR < 0.05) within each 100kb bin were taken. For both gene density and log_2_ fold change, values for genes that overlap the boundary between two bins are counted in both bins.

### Continuous wavelet transformation

Continuous wavelet transformation (CWT), using the transformation function defined in (1) was applied to the binned B-to-A compartment switching, gene density, and log_2_ fold change gene expression values using the *WMTSA* library in R (Percival and Walden, 2000).

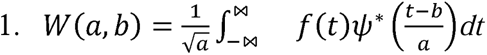

In (1), *W*(*a, b*) is the wavelet transformation of a function *f*(*t*) at a scale *a* ∈ *R*^+*^ and translation value *b* ∈ *R*. Here, *a i*s the scaling parameter, which affects the wavelet’s width. A larger *a* results in a stretched (dilated) wavelet, while a smaller *a* leads to a compressed (contracted) wavelet. *b* is the translation parameter, sliding the wavelet along the chromosome, allowing it to focus on different parts of the chromosome. The function *f*(*t*) is the function that describes the frequency of a signal, e.g., gene density, in chromosomal bin *t*. *Ψ* * is a continuous function in both chromosomal position and frequency and is the complex conjugate of the mother wavelet. In this application, we use a Ricker wavelet. The resulting transformation provides a continuous frequency signal across many scales at their approximate chromosomal locations which can be visualized as a heatmap where the y-axis represents different scales, the x-axis the position along the chromosomal location, and a color gradient is used to visualize the transformed signal frequency. As such, the wavelet transform uses this equation to decompose a signal into components that vary in scale and position, enabling detailed analysis of the (epi)genetic and transcriptomic signal’s characteristics at different resolutions.

### TAD strength analysis

In replicate merged Hi-C maps, Topologically Associated Domains (TADs) were annotated using the arrowhead function from the Juicer tools (Durand et al., 2016). Knight-Ruiz balanced (Knight and Ruiz, 2013) Hi-C maps at 10 kb were used as input and arrowhead function was evoked by adding the “-r 10000”, “-k KR” and “-m 5000” parameters. TAD strength analysis was performed on a combined list of unique TADs identified from control and treated cells per timepoint. Aggregate contact frequency (log_2_ observed/control, at 10 kb resolution) within these combined scaled TADs ± same-sized region was plotted separately for control and treated cells using the fanc aggregate function from the FAN-C package (Kruse et al., 2020). Further, -tad strength and -tads flags were included while calling fanc aggregate function to quantify strength of each TAD in replicate merged control and treated samples separately. To find the contact strength at differentially expressed genes, fanc aggregate function was employed on a bed file of all significant (adjusted p-value <0.05) upregulated (log_2_ fold change >0) and downregulated (log_2_ fold change <0) genes with -tad strength and -tads flags. HiC map file at 10 kb resolution was used as input.

### TAD boundary insulation score analysis

TAD insulation scores across replicate merged control and treated HiC maps were computed using the fanc insulation function from the FAN-C package. HiC map files at 5 kb resolution were used as input and insulation score was computed in 500 kb sliding windows across the genome. Based on this insulation score, we then annotated intervals with insulation score minima as TAD boundaries using fanc boundaries function. Timepoint-specific control and treated TAD boundary coordinates were then combined and the insulation score profiles at and around (±500 kb) the midpoints of TAD boundaries were plotted using the deepTools (Ramirez et al., 2016) plotProfile function.

## Acknowledgements

This material is based upon work supported by the U.S. Department of Energy, Office of Science, through the Biological and Environmental Research (BER) and the Advanced Scientific Computing Research (ASCR) programs under contract number 89233218CNA000001 to Los Alamos National Laboratory (Triad National Security, LLC) (CRS and SS). This work was further supported by the Laboratory Directed Research and Development program of Los Alamos National Laboratory under project numbers 20210134ER (CRS and KYS) and 20210082DR (KYS and SS). This research used resources of the Oak Ridge Leadership Computing Facility at the Oak Ridge National Laboratory, which is supported by the Office of Science of the U.S. Department of Energy under Contract No. DE-AC05-00OR22725. This manuscript has been coauthored by UT-Battelle, LLC under Contract No. DE-AC05-00OR22725 with the U.S. Department of Energy. The United States Government retains and the publisher, by accepting the article for publication, acknowledges that the United States Government retains a non-exclusive, paid-up, irrevocable, world-wide license to publish or reproduce the published form of this manuscript, or allow others to do so, for United States Government purposes. The Department of Energy will provide public access to these results of federally sponsored research in accordance with the DOE Public Access Plan (http://energy.gov/downloads/doe-public-access-plan).

## Author contributions

CRS conceived the research; CRS and VV designed the drug treatment experiments, gene expression, chromatin architecture, accessibility, and epigenetic profiling assays. SA performed the drug treatments, histone ELISA, and microscopy. SA, VV, and ES performed the cell harvest and sample collection. VV performed the sample processing and preparation of ATAC-seq and Hi-C libraries. ES performed the sample processing and preparation of all ChIP-seq libraries. SA performed the RNA extractions and the LANL Genome center prepared RNA-seq libraries. All sequencing was performed at the LANL Genome Center. ES aligned and processed the RNA-seq data. CR aligned and processed the Hi-C and ChIP sequencing data. VV aligned and processed the ATAC-seq data. CR and VV analyzed the Hi-C data. VV integrated all data sets and interpreted the results (Hi-C, ATAC-seq, RNA-seq, and ChIP-seq). Custom python script for comparing profile curves were produced by AHCV. CWT analysis was conceived of by DJ and performed and written about by AHCV, KAS, and CA, and DJ. VV and CRS performed analyses and biological interpretation of all datasets and wrote the manuscript. CRS, KYS, SS, and DJ provided funding for the project.

## References

Andrews, S. (2007). SeqMonk.

Andrews, S. (2010). FastQC: A quality control tool for high throughput sequence data.

Bannister, A.J., and Kouzarides, T. (2011). Regulation of chromatin by histone modifications. Cell Res 21, 381–395.

Baumann, C., Zhang, X.Y., Zhu, L., Fan, Y.H., and De La Fuente, R. (2021). Changes in chromatin accessibility landscape and histone H3 core acetylation during valproic acid-induced differentiation of embryonic stem cells. Epigenet Chromatin 14.

Baylin, S.B. (2005). DNA methylation and gene silencing in cancer. Nat Clin Pract Oncol 2 *Suppl 1*, S4–11.

Beckers, T., Burkhardt, C., Wieland, H., Gimmnich, P., Ciossek, T., Maier, T., and Sanders, K. (2007). Distinct pharmacological properties of second generation HDAC inhibitors with the benzamide or hydroxamate head group. Int J Cancer 121, 1138–1148.

Caslini, C., Hong, S., Ban, Y.J., Chen, X.S., and Ince, T.A. (2019). HDAC7 regulates histone 3 lysine 27 acetylation and transcriptional activity at super-enhancer-associated genes in breast cancer stem cells. Oncogene 38, 6599–6614.

Chen, S., Zhou, Y., Chen, Y., and Gu, J. (2018). fastp: an ultra-fast all-in-one FASTQ preprocessor. Bioinformatics 34, i884–i890.

Chuang, D.M., Leng, Y., Marinova, Z., Kim, H.J., and Chiu, C.T. (2009). Multiple roles of HDAC inhibition in neurodegenerative conditions. Trends Neurosci 32, 591–601.

Dang, Y., Li, S., Zhao, P., Xiao, L., Wang, L., Shi, Y., Luo, L., Wang, S., Wang, H., and Zhang, K. (2022). The lysine deacetylase activity of histone deacetylases 1 and 2 is required to safeguard zygotic genome activation in mice and cattle. Development 149.

Dixon, J.R., Selvaraj, S., Yue, F., Kim, A., Li, Y., Shen, Y., Hu, M., Liu, J.S., and Ren, B. (2012). Topological domains in mammalian genomes identified by analysis of chromatin interactions. Nature 485, 376–380.

Dobin, A., Davis, C.A., Schlesinger, F., Drenkow, J., Zaleski, C., Jha, S., Batut, P., Chaisson, M., and Gingeras, T.R. (2013). STAR: ultrafast universal RNA-seq aligner. Bioinformatics 29, 15–21.

Durand, N.C., Shamim, M.S., Machol, I., Rao, S.S., Huntley, M.H., Lander, E.S., and Aiden, E.L. (2016). Juicer Provides a One-Click System for Analyzing Loop-Resolution Hi-C Experiments. Cell Syst 3, 95–98.

Faust, G.G., and Hall, I.M. (2014). SAMBLASTER: fast duplicate marking and structural variant read extraction. Bioinformatics 30, 2503–2505.

Finn, E.H., Pegoraro, G., Brandao, H.B., Valton, A.L., Oomen, M.E., Dekker, J., Mirny, L., and Misteli, T. (2019). Extensive Heterogeneity and Intrinsic Variation in Spatial Genome Organization. Cell 176, 1502–1515 e1510.

Gao, T.S., and Qian, J. (2020). EnhancerAtlas 2.0: an updated resource with enhancer annotation in 586 tissue/cell types across nine species. Nucleic Acids Research 48, D58–D64.

Gaspar, J.M. (2018). Improved peak-calling with MACS2. bioRxiv.

Glozak, M.A., and Seto, E. (2007). Histone deacetylases and cancer. Oncogene 26, 5420–5432.

Halsall, J.A., Turan, N., Wiersma, M., and Turner, B.M. (2015). Cells adapt to the epigenomic disruption caused by histone deacetylase inhibitors through a coordinated, chromatin-mediated transcriptional response. Epigenet Chromatin 8.

Harris, H.L., Gu, H., Olshansky, M., Wang, A., Farabella, I., Eliaz, Y., Kalluchi, A., Krishna, A., Jacobs, M., Cauer, G., et al. (2023). Chromatin alternates between A and B compartments at kilobase scale for subgenic organization. Nature Communications 14.

Hnisz, D., Shrinivas, K., Young, R.A., Chakraborty, A.K., and Sharp, P.A. (2017). A Phase Separation Model for Transcriptional Control. Cell 169, 13–23.

Hogg, S.J., Motorna, O., Cluse, L.A., Johanson, T.M., Coughlan, H.D., Raviram, R., Myers, R.M., Costacurta, M., Todorovski, I., Pijpers, L., et al. (2021). Targeting histone acetylation dynamics and oncogenic transcription by catalytic P300/CBP inhibition. Mol Cell 81, 2183–2200 e2113.

Jacques Dainat, D.H., Dr.K. D. Murray, Ed Davis, Kathryn Crouch, LucileSol, Nuno Agostinho, pascal-git, Zachary Zollman, & tayyrov (2023). NBISweden/AGAT: AGAT-v1.2.0 (v1.2.0). Zenodo.

Keppler, B.R., and Archer, T.K. (2008). Chromatin-modifying enzymes as therapeutic targets--Part 1. Expert Opin Ther Targets 12, 1301–1312.

Knight, P.A., and Ruiz, D. (2013). A fast algorithm for matrix balancing. Ima J Numer Anal 33, 1029–1047.

Kruse, K., Hug, C.B., and Vaquerizas, J.M. (2020). FAN-C: a feature-rich framework for the analysis and visualisation of chromosome conformation capture data. Genome Biol 21, 303.

Lai, W.K.M., and Pugh, B.F. (2017). Understanding nucleosome dynamics and their links to gene expression and DNA replication. Nat Rev Mol Cell Biol 18, 548–562.

Li, H., and Durbin, R. (2009). Fast and accurate short read alignment with Burrows-Wheeler transform. Bioinformatics 25, 1754–1760.

Li, Y., and Seto, E. (2016). HDACs and HDAC Inhibitors in Cancer Development and Therapy. Cold Spring Harb Perspect Med 6.

Love, M.I., Huber, W., and Anders, S. (2014). Moderated estimation of fold change and dispersion for RNA-seq data with DESeq2. Genome Biol 15, 550.

Mansisidor, A.R., and Risca, V.I. (2022). Chromatin accessibility: methods, mechanisms, and biological insights. Nucleus 13, 236–276.

Mirny, L.A., Imakaev, M., and Abdennur, N. (2019). Two major mechanisms of chromosome organization. Curr Opin Cell Biol 58, 142–152.

Misteli, T. (2020). The Self-Organizing Genome: Principles of Genome Architecture and Function. Cell 183, 28–45.

Morgan, M.A., and Shilatifard, A. (2015). Chromatin signatures of cancer. Genes Dev 29, 238–249.

Myers, R.M., Brown, M., Li, W., et al. (2008). Model-based analysis of ChIP-Seq (MACS). Genome Biol 9, R137.

Nurk, S., Koren, S., Rhie, A., Rautiainen, M., Bzikadze, A.V., Mikheenko, A., Vollger, M.R., Altemose, N., Uralsky, L., Gershman, A., et al. (2022). The complete sequence of a human genome. Science 376, 44–53.

Percival, D.B., and Walden, A.T. (2000). Wavelet methods for time series analysis (Cambridge; New York: Cambridge University Press).

Putri, G.H., Anders, S., Pyl, P.T., Pimanda, J.E., and Zanini, F. (2022). Analysing high-throughput sequencing data in Python with HTSeq 2.0. Bioinformatics 38, 2943–2945.

Quinlan, A.R., and Hall, I.M. (2010). BEDTools: a flexible suite of utilities for comparing genomic features. Bioinformatics 26, 841–842.

Ramirez, F., Ryan, D.P., Gruning, B., Bhardwaj, V., Kilpert, F., Richter, A.S., Heyne, S., Dundar, F., and Manke, T. (2016). deepTools2: a next generation web server for deep-sequencing data analysis. Nucleic Acids Res 44, W160–165.

Rao, S.S., Huntley, M.H., Durand, N.C., Stamenova, E.K., Bochkov, I.D., Robinson, J.T., Sanborn, A.L., Machol, I., Omer, A.D., Lander, E.S., et al. (2014). A 3D map of the human genome at kilobase resolution reveals principles of chromatin looping. Cell 159, 1665–1680.

Reed, K.S.M., Davis, E.S., Bond, M.L., Cabrera, A., Thulson, E., Quiroga, I.Y., Cassel, S., Woolery, K.T., Hilton, I., Won, H., et al. (2022). Temporal analysis suggests a reciprocal relationship between 3D chromatin structure and transcription. Cell Rep 41, 111567.

Roth, C., Venu, V., Job, V., Lubbers, N., Sanbonmatsu, K.Y., Steadman, C.R., and Starkenburg, S.R. (2023). Improved quality metrics for association and reproducibility in chromatin accessibility data using mutual information. BMC Bioinformatics 24, 441.

Sanborn, A.L., Rao, S.S., Huang, S.C., Durand, N.C., Huntley, M.H., Jewett, A.I., Bochkov, I.D., Chinnappan, D., Cutkosky, A., Li, J., et al. (2015). Chromatin extrusion explains key features of loop and domain formation in wild-type and engineered genomes. Proc Natl Acad Sci U S A 112, E6456–6465.

Sanchez, G.J., Richmond, P.A., Bunker, E.N., Karman, S.S., Azofeifa, J., Garnett, A.T., Xu, Q., Wheeler, G.E., Toomey, C.M., Zhang, Q., et al. (2018). Genome-wide dose-dependent inhibition of histone deacetylases studies reveal their roles in enhancer remodeling and suppression of oncogenic super-enhancers. Nucleic Acids Res 46, 1756–1776.

Schmitt, A. D., Hu, M., Jung, I., Xu, Z., Qiu, Y., Tan, C. L., … Ren, B. (2016). A Compendium of Chromatin Contact Maps Reveals Spatially Active Regions in the Human Genome. Cell Rep, 17(8), 2042–2059. doi:10.1016/j.celrep.2016.10.061

Schoenfelder, S., and Fraser, P. (2019). Long-range enhancer-promoter contacts in gene expression control. Nat Rev Genet 20, 437–455.

Slaughter, M.J., Shanle, E.K., Khan, A., Chua, K.F., Hong, T., Boxer, L.D., Allis, C.D., Josefowicz, S.Z., Garcia, B.A., Rothbart, S.B., et al. (2021). HDAC inhibition results in widespread alteration of the histone acetylation landscape and BRD4 targeting to gene bodies. Cell Rep 34, 108638.

Venu, V., Roth, C., Adikari, S.H., Small, E.M., Starkenburg, S.R., Sanbonmatsu, K.Y., and Steadman, C.R. (2023). Vaccinia virus infection induces concurrent alterations in host chromatin architecture, accessibility, and gene expression. bioRxiv.

Wang, R., Lee, J.H., Kim, J., Xiong, F., Hasani, L.A., Shi, Y., Simpson, E.N., Zhu, X., Chen, Y.T., Shivshankar, P., et al. (2023). SARS-CoV-2 restructures host chromatin architecture. Nat Microbiol 8, 679–694.

Wolffe, A.P. (2001). Chromatin remodeling: why it is important in cancer. Oncogene 20, 2988–2990.

Wu, Q., Cheng, Z.Y., Zhu, J., Xu, W.Q., Peng, X.J., Chen, C.B., Li, W.T., Wang, F.S., Cao, L.J., Yi, X.L., et al. (2015). Suberoylanilide Hydroxamic Acid Treatment Reveals Crosstalks among Proteome, Ubiquitylome and Acetylome in Non-Small Cell Lung Cancer A549 Cell Line. Sci Rep-Uk 5.

Yan, K.K., Yardimci, G.G., Yan, C., Noble, W.S., and Gerstein, M. (2017). HiC-spector: a matrix library for spectral and reproducibility analysis of Hi-C contact maps. Bioinformatics 33, 2199–2201.

Zahir, F.R., and Brown, C.J. (2011). Epigenetic impacts on neurodevelopment: pathophysiological mechanisms and genetic modes of action. Pediatr Res 69, 92R–100R.

Zhang, Y., Liu, T., Meyer, C.A., Eeckhoute, J., Johnson, D.S., Bernstein, B.E., Nusbaum, C., Zhang, Y., See, Y. X., Tergaonkar, V., & Fullwood, M. J. (2022). Long-Distance Repression by Human Silencers: Chromatin Interactions and Phase Separation in Silencers. Cells, 11(9). doi:10.3390/cells11091560

