## Supplementary_Material for "Epigenomic manipulation reveals the relationship between locus specific chromatin dynamics and gene expression"

for

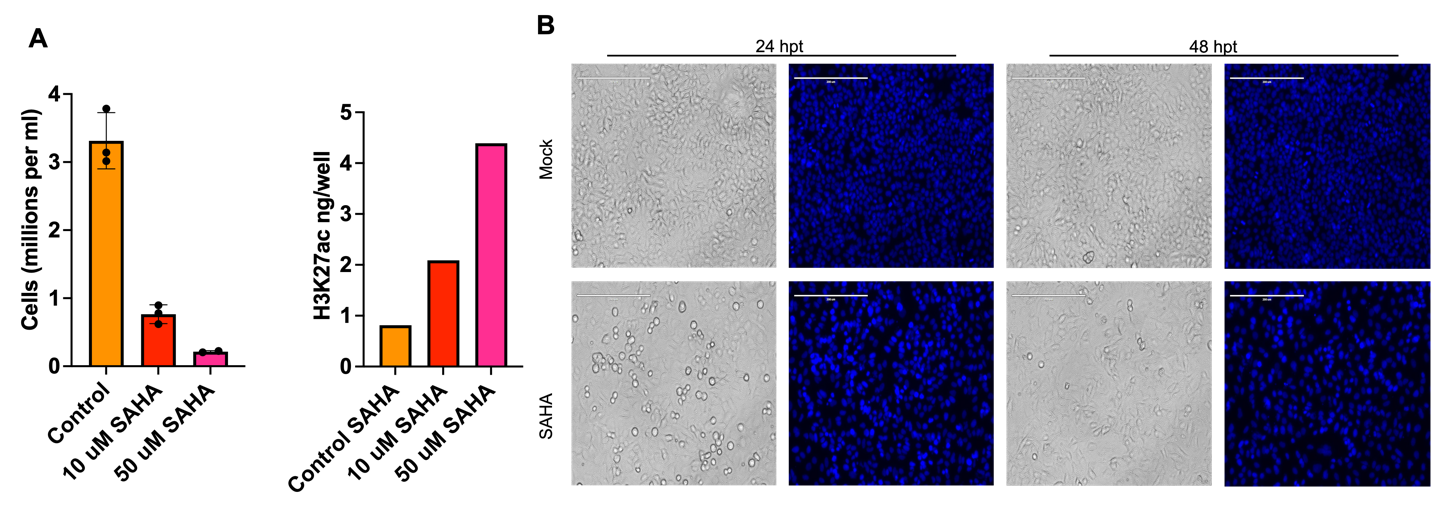

***Figure S 1:*** *A) Cell counts (left) and global H3K27ac measurements via ELISA (right) at 48 hours post treatment (hpt) demonstrate 10 μM SAHA induces considerable changes in acetylation while retaining viable cells compared to 50 μM SAHA. B) Bright-field and DAPI stained images of control (Mock-treated) and 10 μM SAHA-treated (SAHA) cells at 24 and 48 hpt are shown at 20X magnification. Scale bar : 200 μm.*

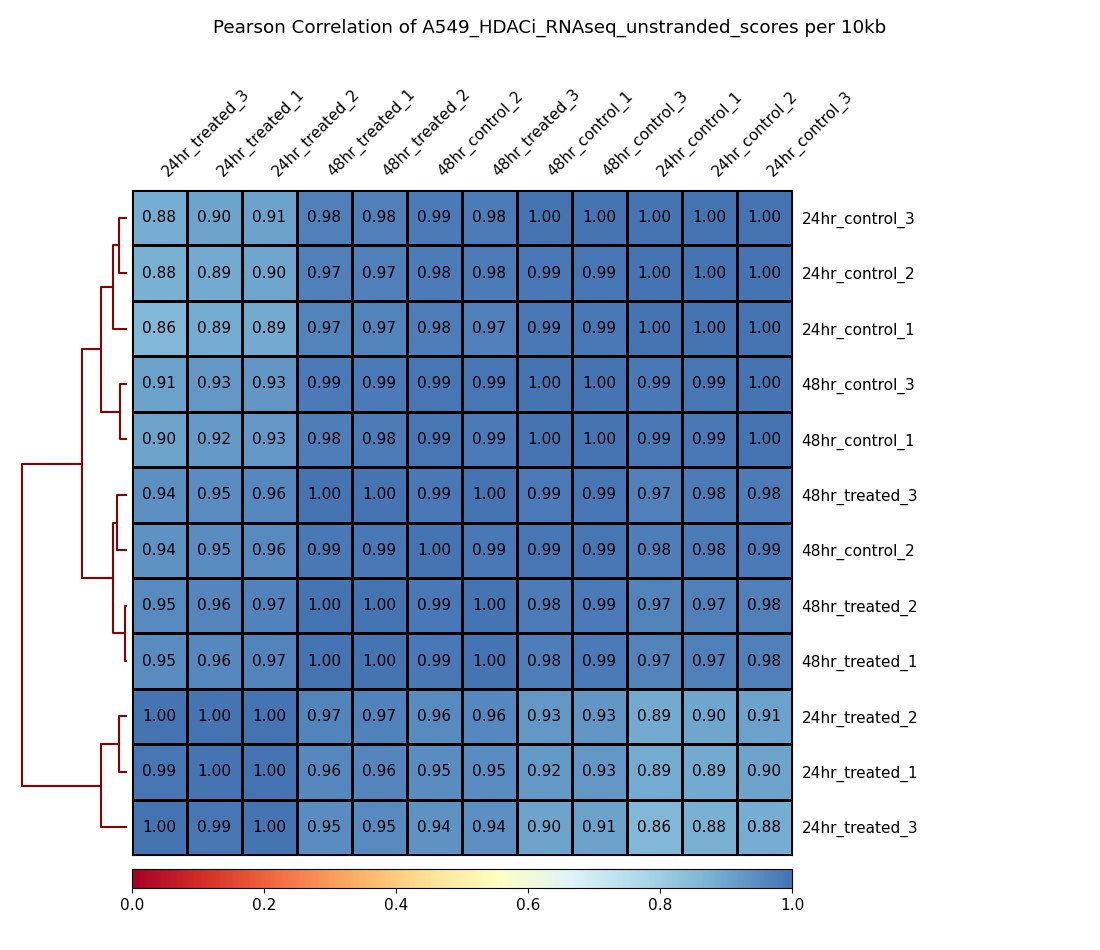

**Figure S 2**: Genome-wide Pearson correlation between RNA-seq samples at 10 kb resolution.

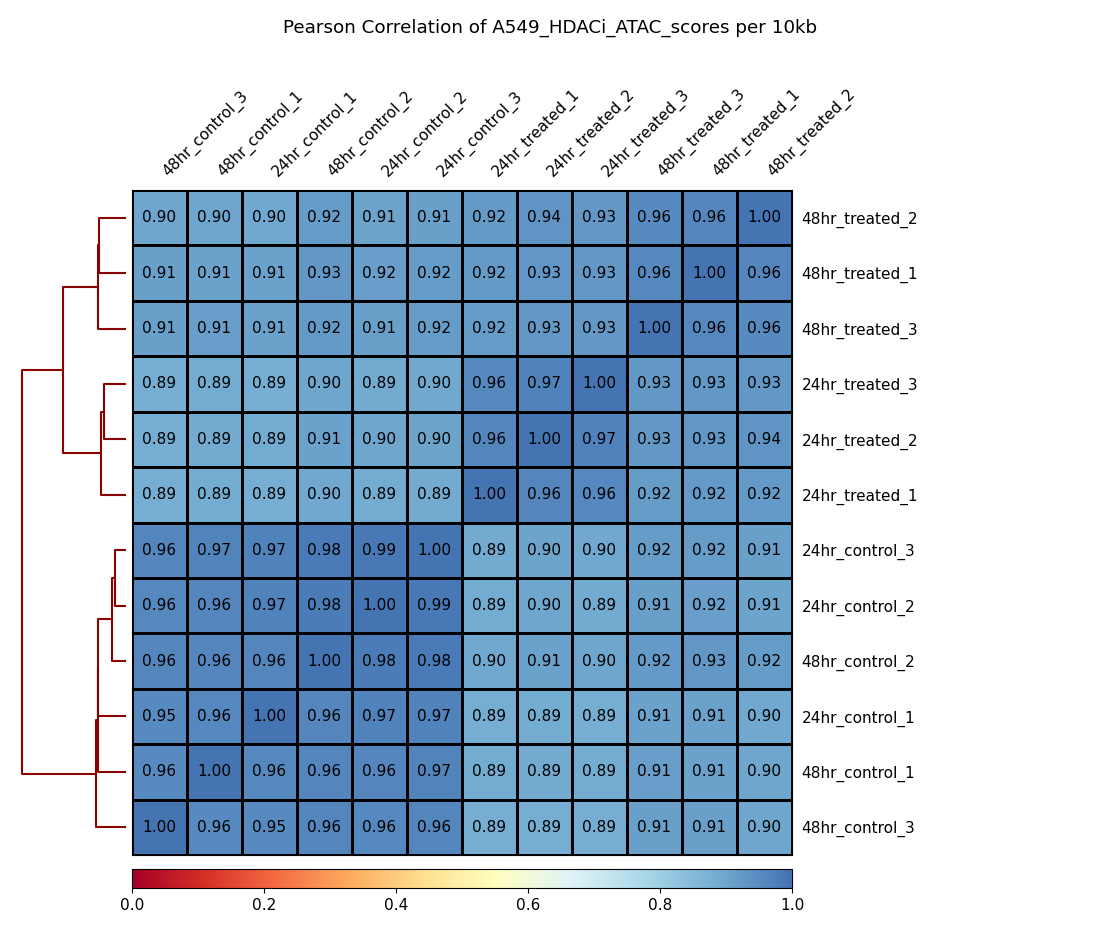

**Figure S 3:** Genome-wide Pearson correlation between ATAC-seq samples at 10 kb resolution.

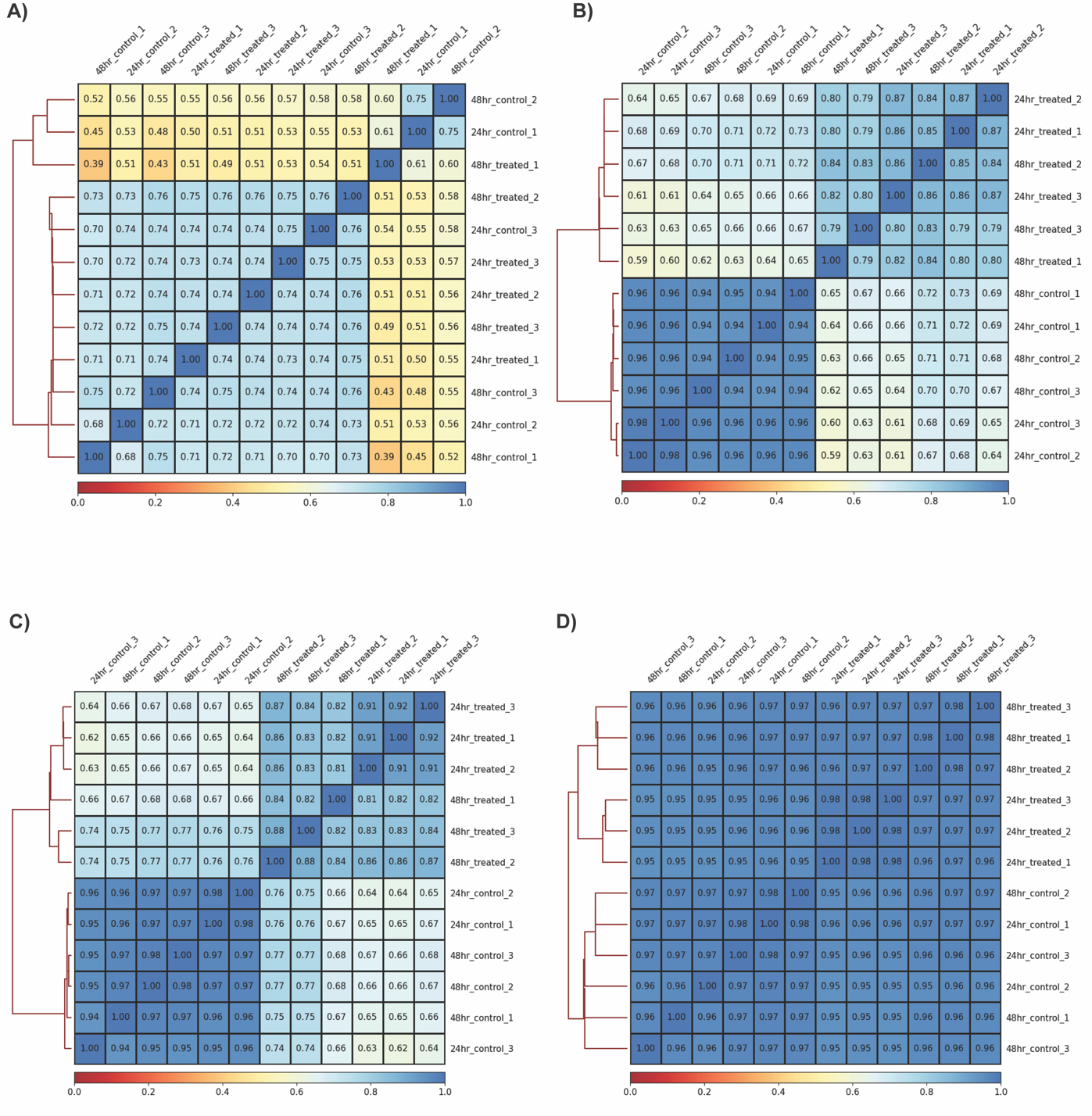

**Figure S 4**: Genome-wide Pearson correlation between A) H3, B) H3ac, C) H3K27ac, and D )H3K4me3 ChIP-seq samples at 10 kb resolution using SPMR normalized reads. Three outlier samples identified with H3-ChIP (48hr_control_2, 48hr_treated_1, 24hr_control_1) were excluded from further analysis.

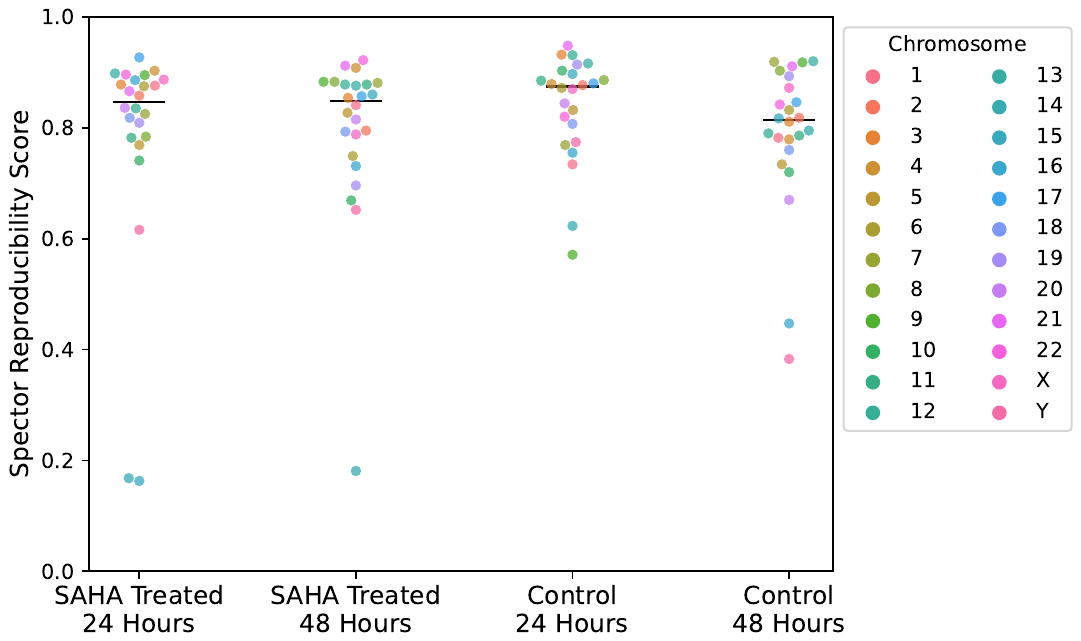

***Figure S 5:*** *Chromosome-wise Spector similarity score estimated for Hi-C data at 500 kb resolution between biological replicates for each condition per time point. Horizontal black lines mark median score value.*

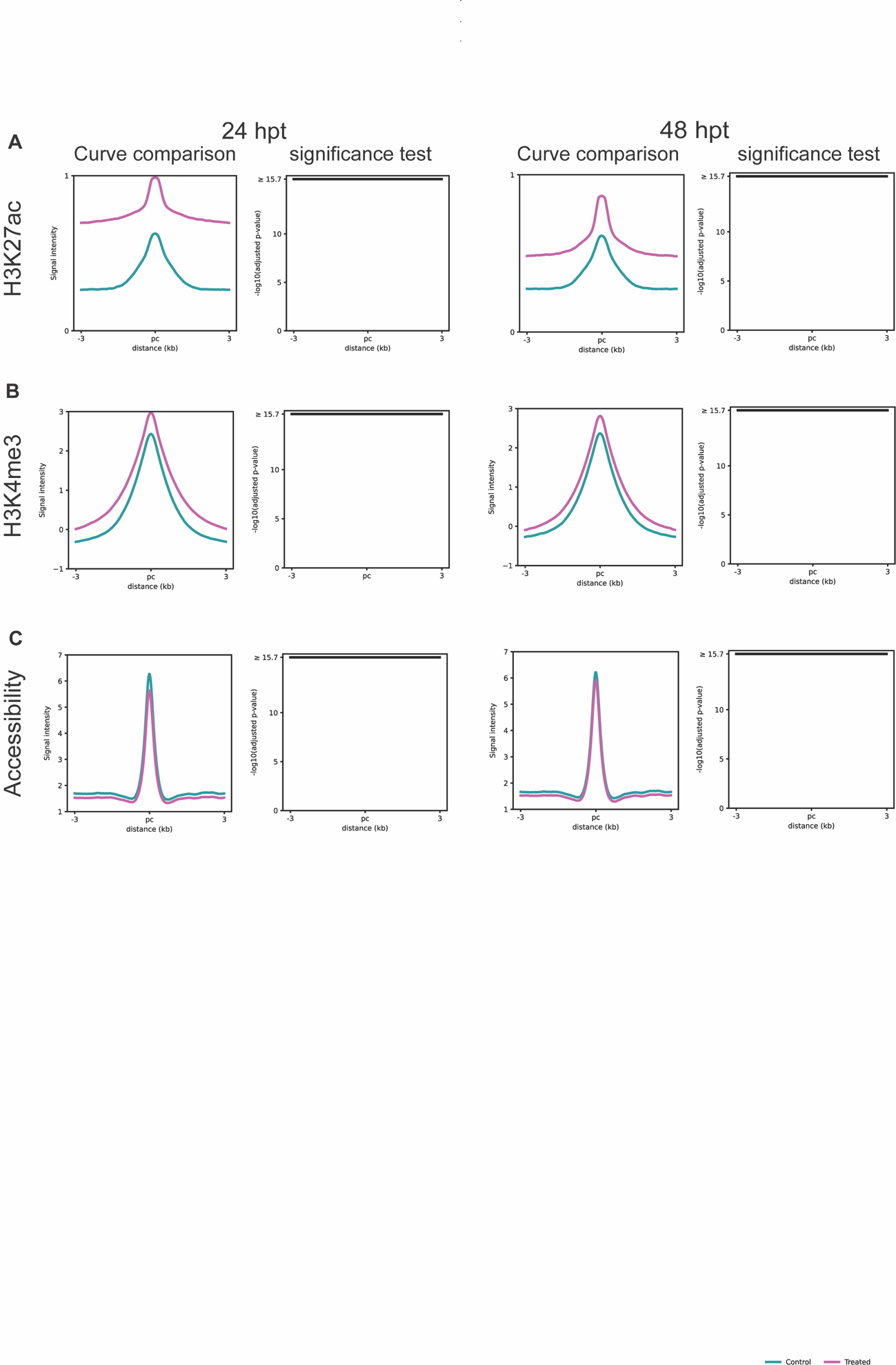

Figure S 6: A) H3K27ac, B) H3K4me3, and C) Chromatin accessibility global profile curve comparison and significance profile (wilcox rank sum test, negative log_2_ adjusted p-value) for control (teal) and treated (pink) at 24 hpt (left) and 48 hpt (right). For all comparisons, negative log_2_ adjusted p-value above >15.7 indicates significant difference between control and treated profiles.

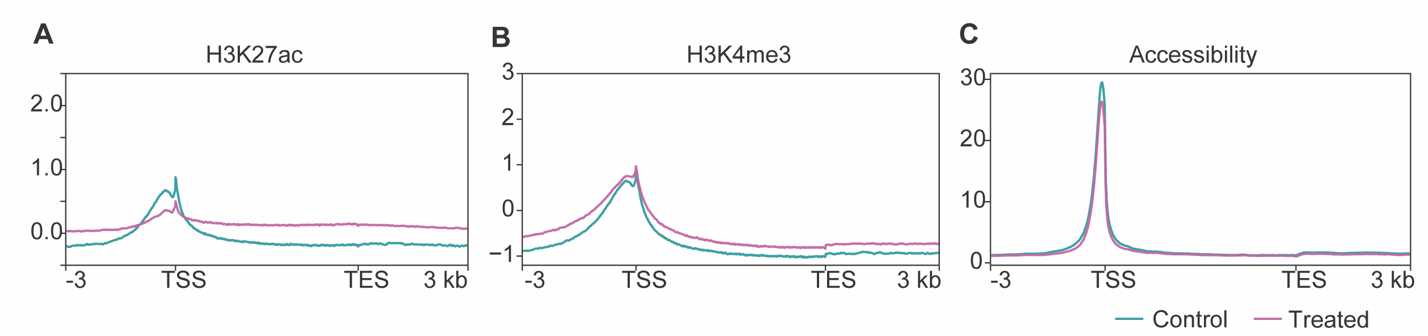

**Figure S 7:** H3 normalized H3K27ac and H3K4me3 profiles and SPMR normalized chromatin accessibility at 24 hpt across all genes. Genes are scaled to 5 kb and gene ± 3 kb region is shown. Control (teal) and treated (pink) profiles are overlaid.

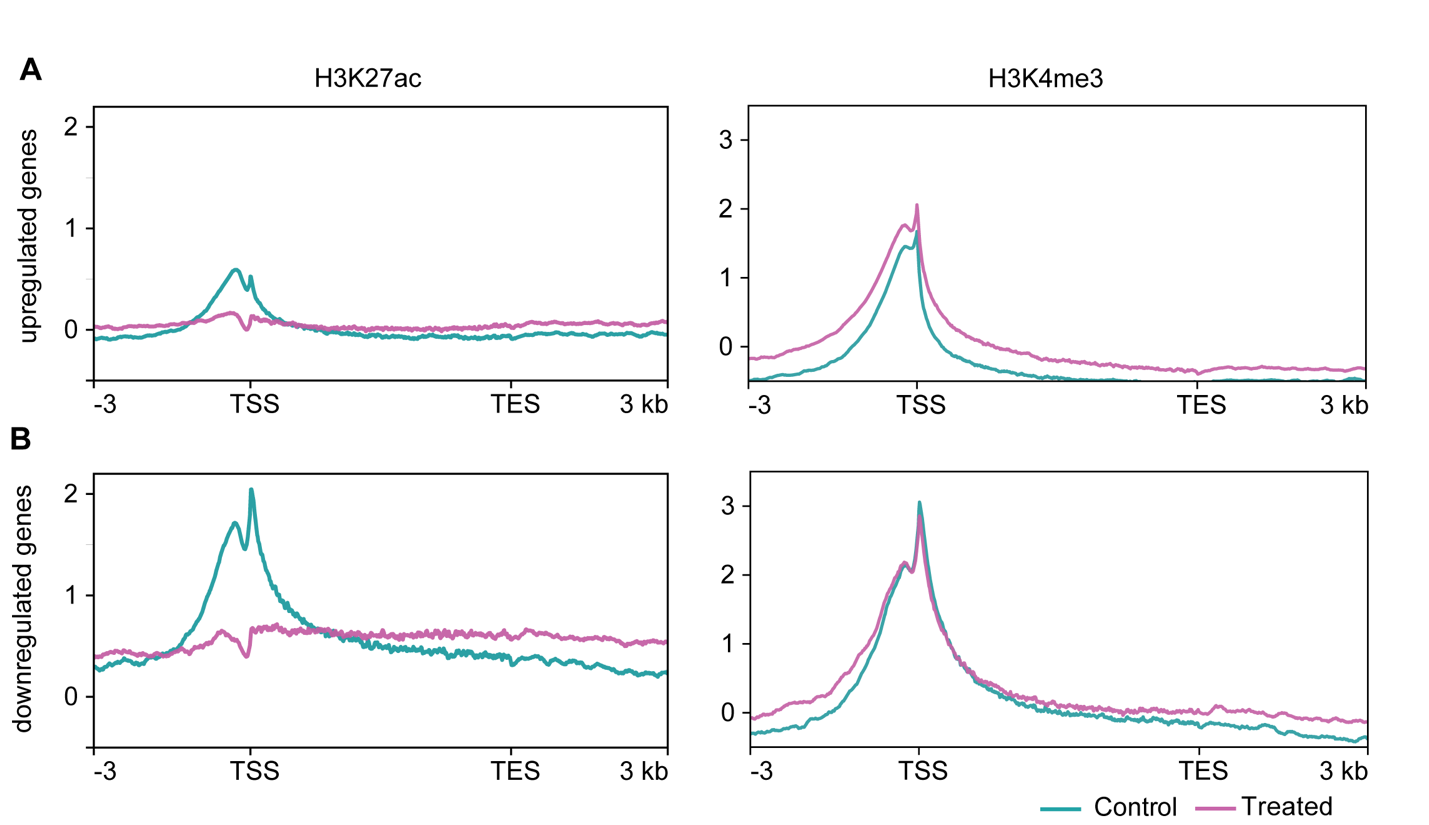

**Figure S 8:** Input normalized H3K27ac (left) and H3K4me3 (right) profiles across 24 hpt (A) upregulated and (B) down regulated genes with greater than 2-fold change. All genes are scaled to 5 kb and gene ± 3 kb region is shown. Control (teal) and treated (pink) profiles are overlaid

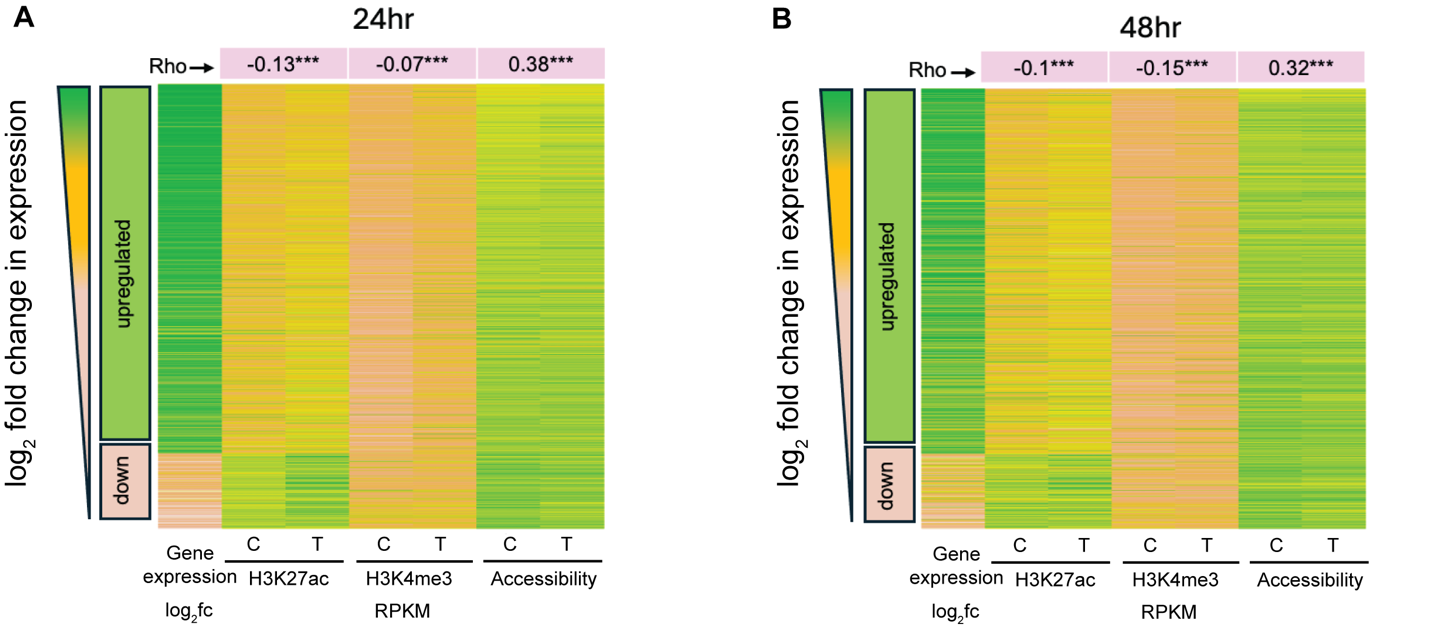

**Figure S 9:** Heatmap representing log2 fold change in gene expression of significantly differentially expressed genes at 24 hpt (p-adj <0.05 and log2f c>2) sorted in the order of positive to negative fold change in expression. Control (C) and treated (T) values of gene body (TSS+500bp till TES) H3K27ac, H3K4me3, and accessibility are shown in the subsequent columns for those genes. Spearman rank correlation (Rho) between fold change in expression vs fold change in each of the features is given on top. (*** denotes p-value <0.001).

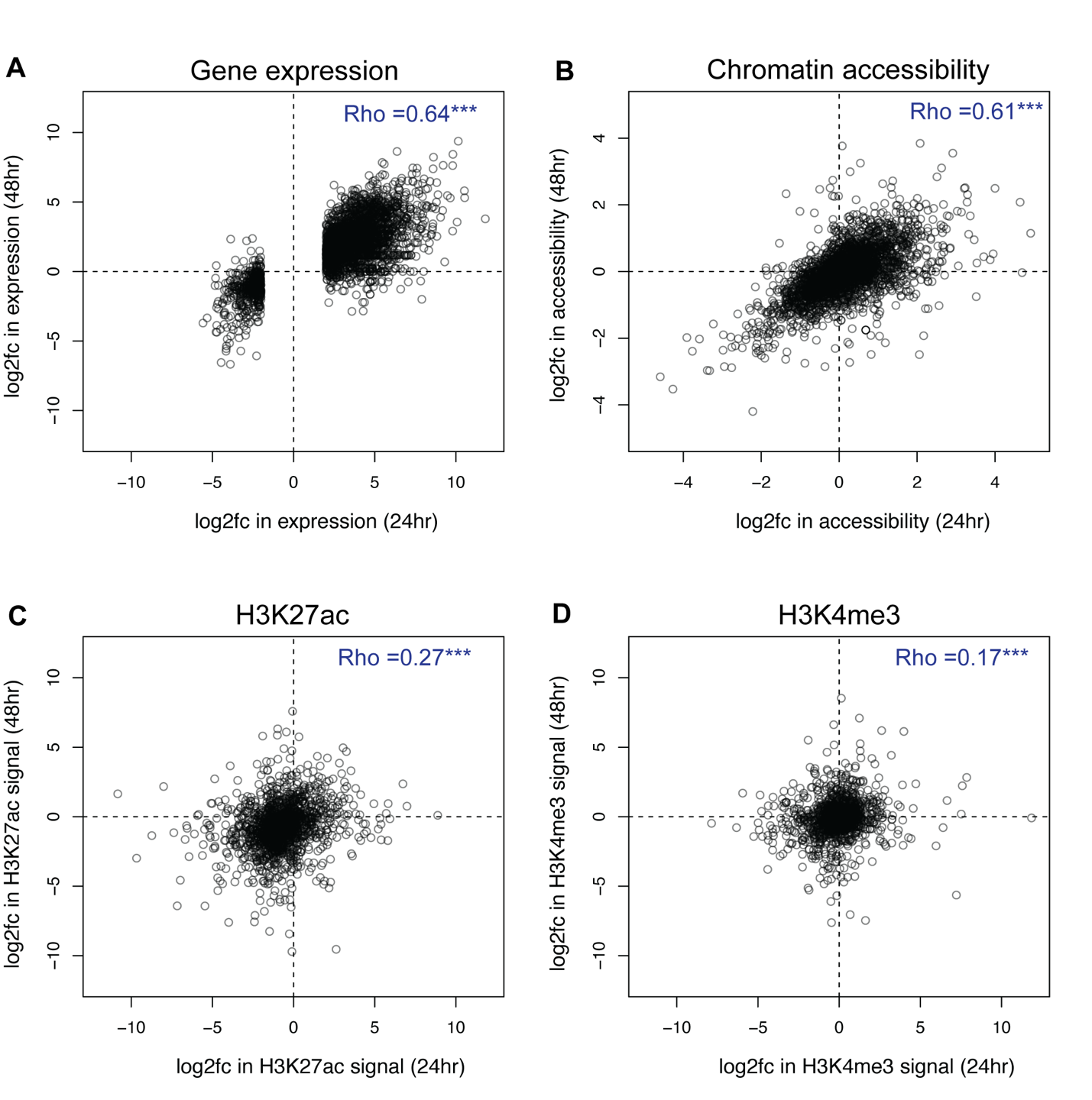

**Figure S 10:** For significant differential genes at 24 hpt (adjusted p-value <0.05 and abs(log2 fold change) >2), fold change in their A) gene expression and fold change in their promoter B) chromatin accessibility, C) H3K27ac, and D) H3K4me3, signal at 24 hpt vs 48 hpt is plotted as a scatter plot. Spearman correlation value is annotated in corresponding plot with indication of statistical significance (*** denotes p-value<0.001).

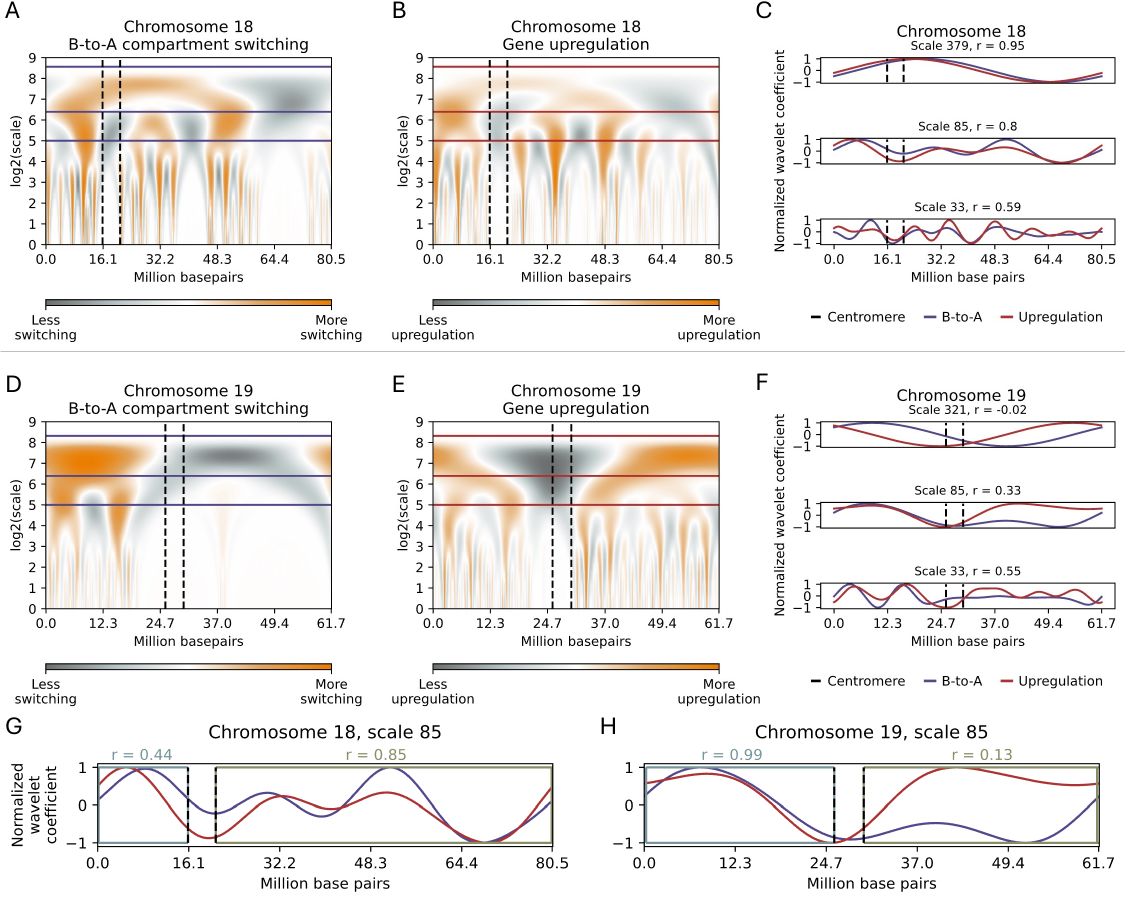

**Figure S 11: Concordance between Hi-C B-to-A compartment switching and gene upregulation differs between gene-rich and gene-poor chromosomes.** A) Density of the Hi-C B-to-A compartment switching signal at 100 kb windows along chromosome 18 using continuous wavelet transformation. B) Density of upregulated genes at 100 kb intervals along chromosome 18 using continuous wavelet transformation. C) The intersection of the B-to-A wavelet and the gene upregulation at different scales on chromosome 18. The Pearson’s correlation coefficient (r) at each reported scale is provided in the title. D) Density of the Hi-C B-to-A compartment switching signal at 100 kb windows along chromosome 19 using continuous wavelet transformation. E) Density of upregulated genes at 100kb intervals along chromosome 19 using continuous wavelet transformation. F) The intersection of the B-to-A wavelet and the gene upregulation at different scales on chromosome 19. The Pearson’s correlation coefficient (r) at each reported scale is provided in the title. G) Correlations between the B-to-A wavelet (Blue) and the upregulation wavelet (Red) at scale 85 on the p arm (annotated in slate blue) and q arm (annotated in olive green) on chromosome 18. H) Correlations between the B-to-A (Blue) wavelet and the upregulation wavelet (Red) at scale 85 on the p arm (annotated in slate blue) and q arm (annotated in olive green) on chromosome 19.

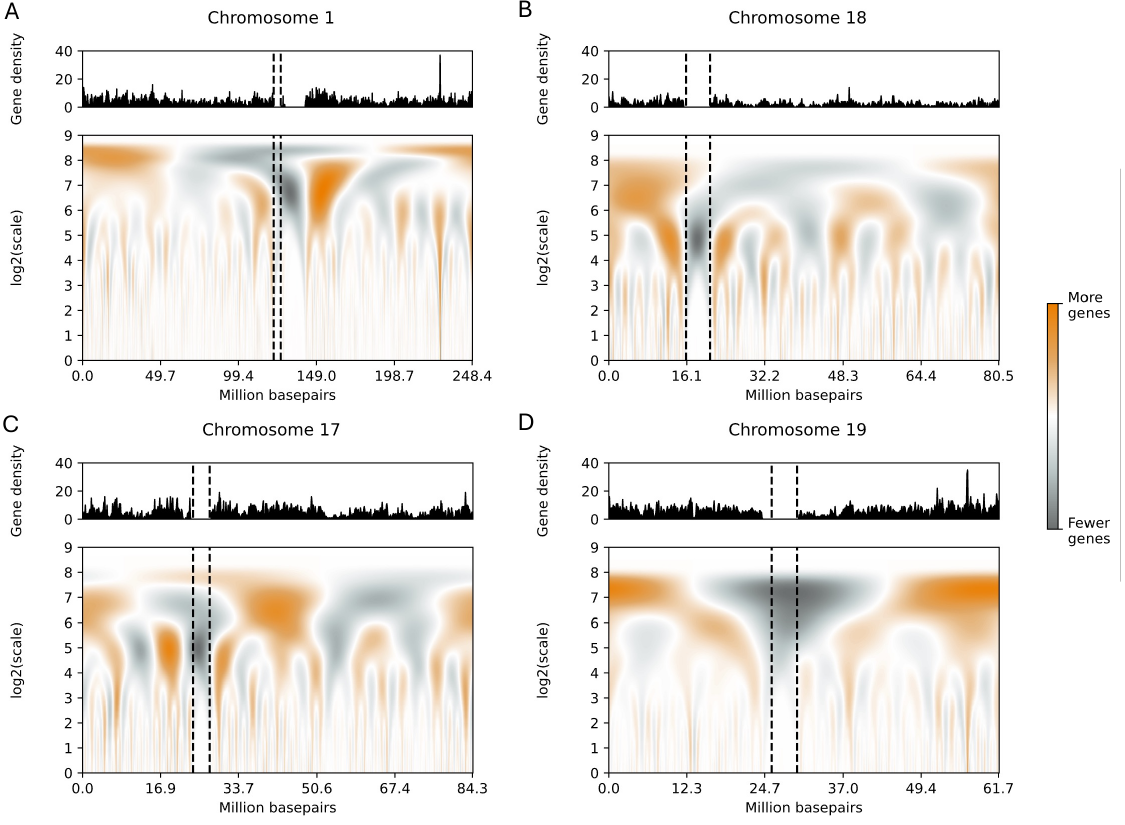

**Figure S 12: Gene density signal top and wavelets.** A) The gene density signal (top) and the continuous wavelet transformation (bottom) of the gene density signal within 100 kb bins along chromosome 1. Panels B, C, and D report the same results for chromosomes 17, 18, and 19, respectively. The dotted lines indicate the location of the centromere.

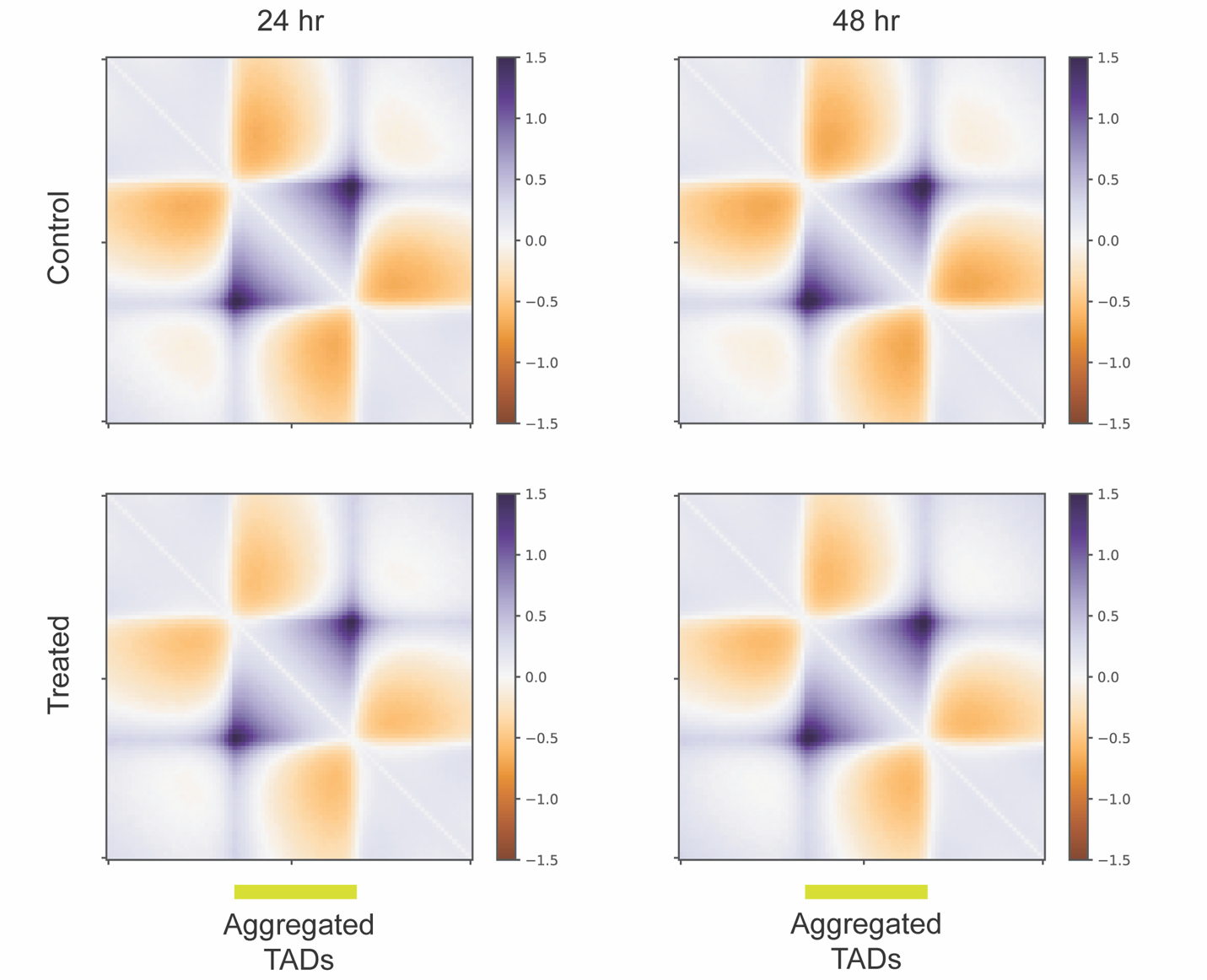

**Figure S 13: Moderate TAD weakening after treatment**. Average log_2_ observed over expected intra-TAD contact frequency within scaled aggregated TADs ± same sized flanking regions are shown for 24 hpt (left) and 48 hpt (right) Control (top) and Treated (bottom) cells.

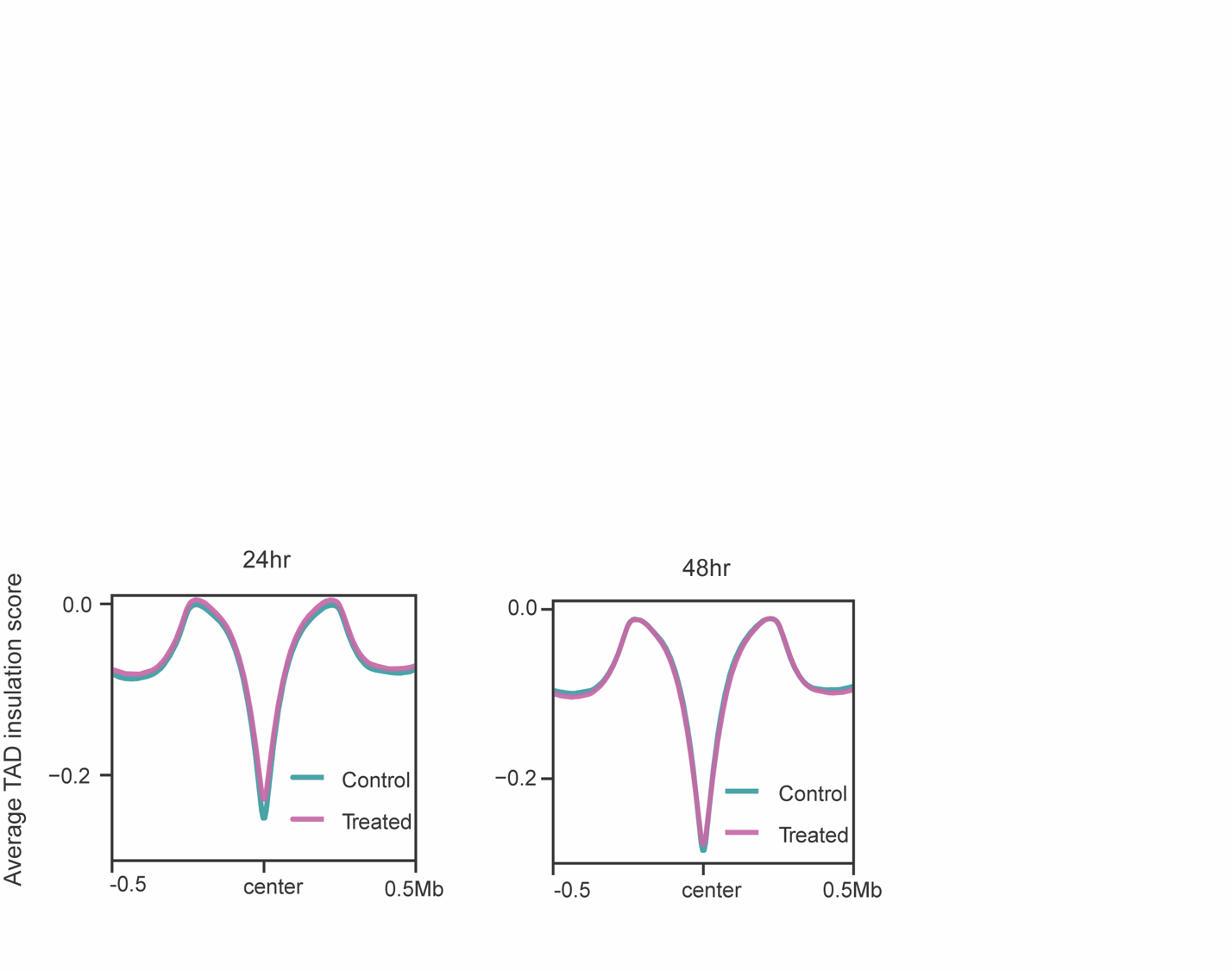

**Figure S 14: TAD boundaries are largely conserved after treatment.** Insulation score computed from all TAD boundary midpoints ± 500 kb region is shown as an average profile plot for 24 hr (left) and 48 hr (right) cells Pink: Treated, Teal: Control.

.

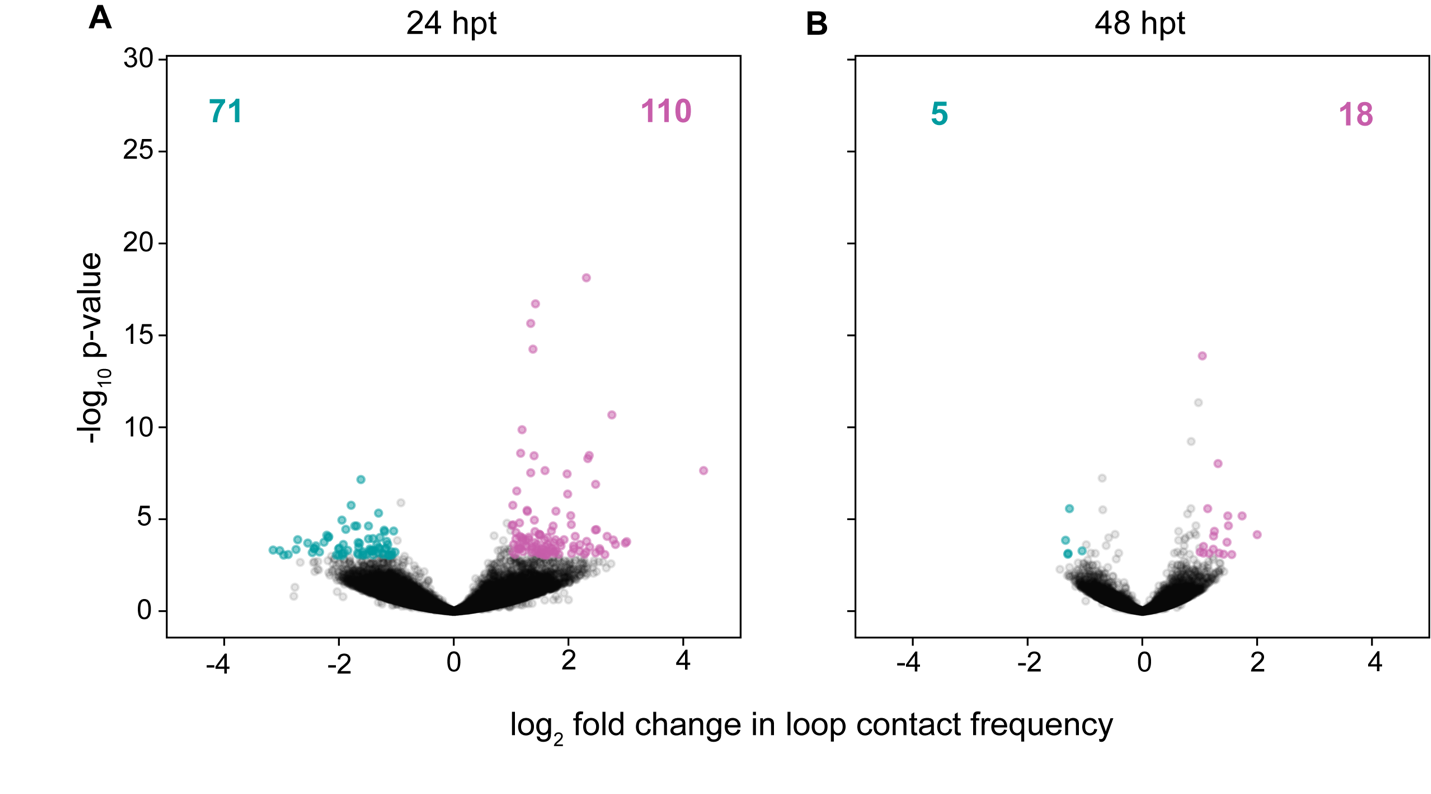

**Figure S 15:** Volcano plots represent loop comparison between control and treated cells in A) 24 hpt and B) 48 hpt. Black dots represent all called loops. Significant control differential loops (adjusted p-value <0.05 and log2 fold change <-2 ) are shown in teal and significant treated loops differential loops (adjusted p-value <0.05 and log2 fold change >2) are shown in pink.

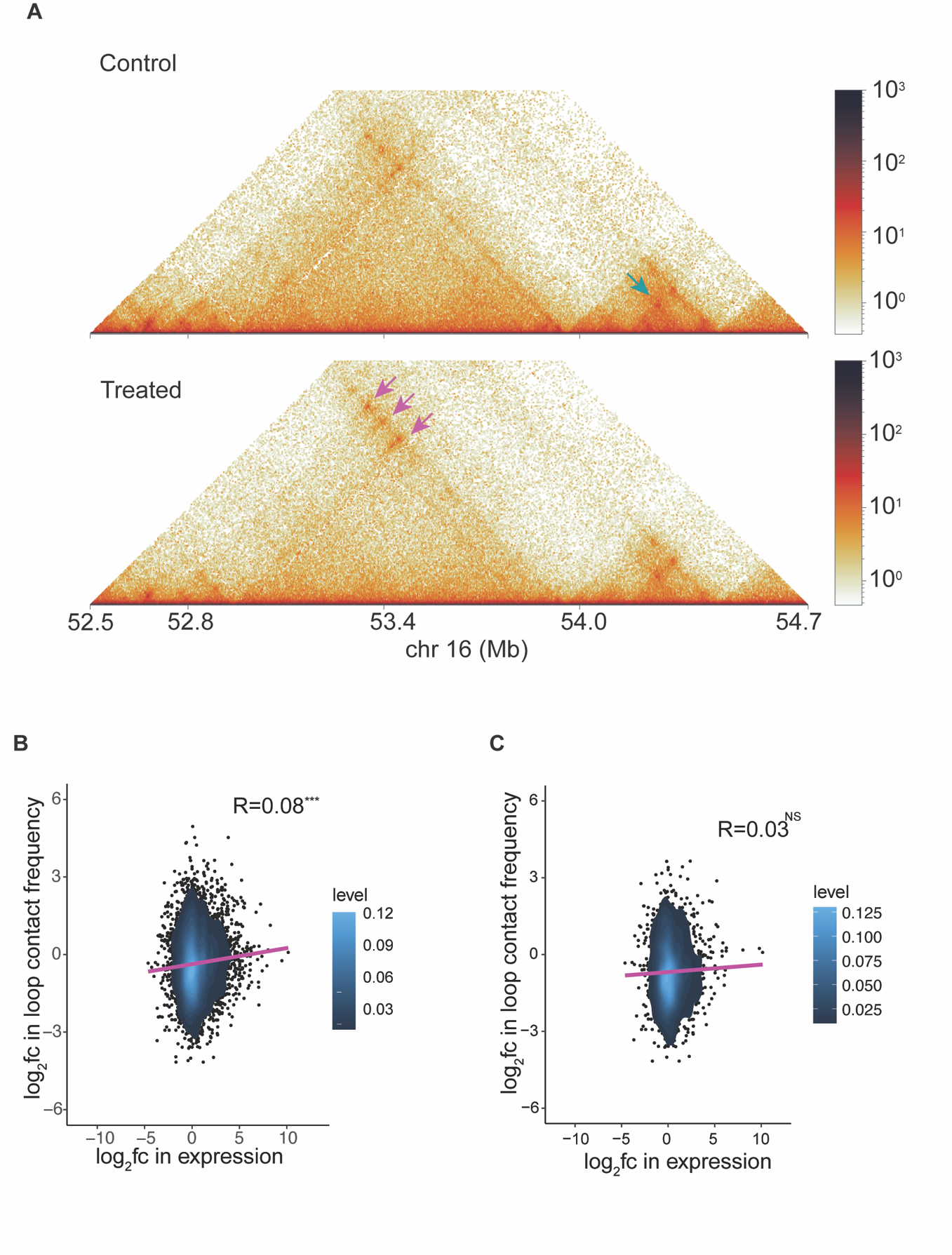

**Figure S 16:At the genome-wide level, loop changes are weak and less predictive of gene expression changes.** A) Control vs Treated Hi-C heatmap from 24 hpt showing an example region comprised of strengthened (marked with pink rectangle) and weakened (marked with teal rectangle) loops after HDACi treatment. B) Scatter plot overlayed with density heatmap presenting the correlation between log_2_ fold change in gene expression and loop strength of promoter overlapping loop anchors. Linear fit is shown by the purple line and the Pearson correlation value (R) and statistical significance (p-value <0.01) are denoted in the plot.C) For genes with an active enhancer promoter loop, their log2 fold change in expression vs log2 fold change in enhancer promoter loop is plotted as a scatter plot with overlaid density heatmap. Linear fit is shown by the purple line; and the Pearson correlation value R is denoted in the plot. The correlation found to be statistically nonsignificant. df = 1660, p-value = 0.166

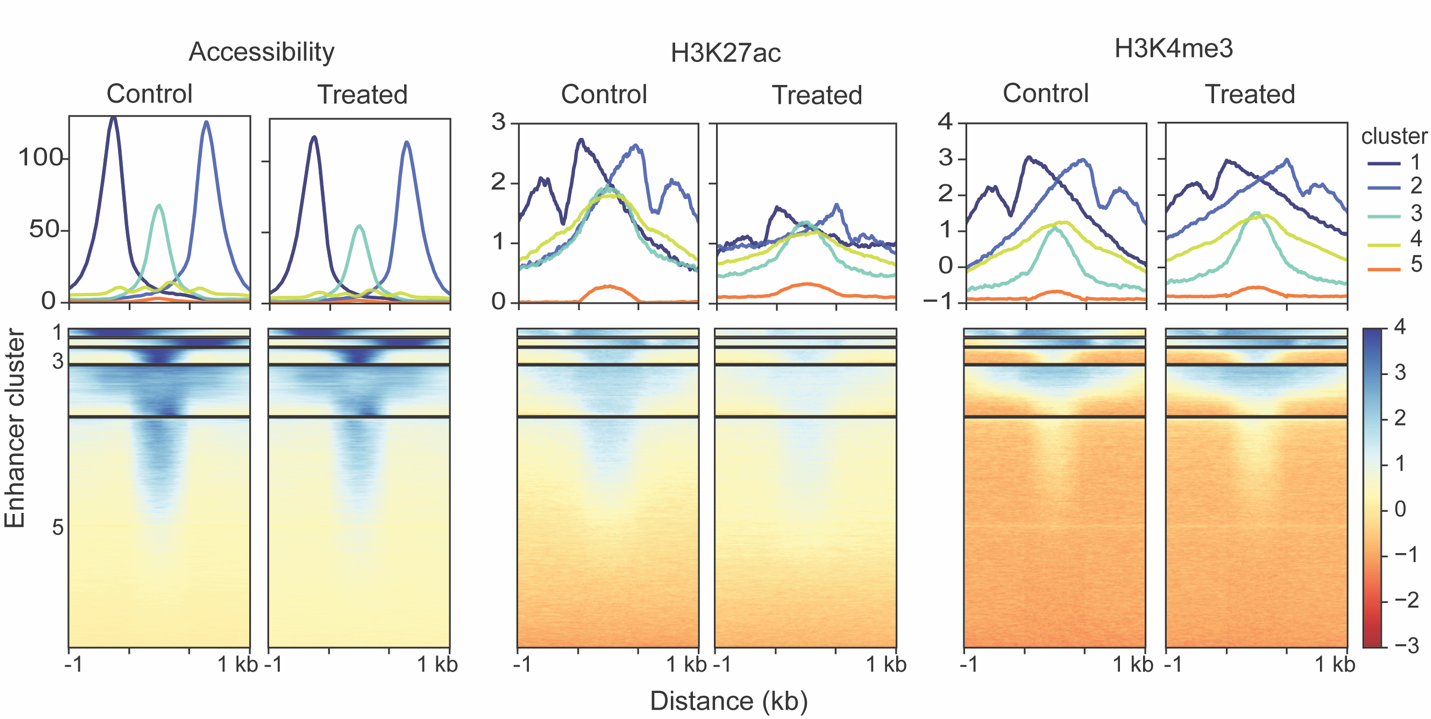

**Figure S 17:**  **A subset of annotated enhancers are accessible and possess active chromatin marks.** Control and treated accessibility, H3K27ac, and H3K4me3 profiles across all annotated enhancers ± 1 kb are shown. A clustering analysis (kmeans) separates enhancers into 5 clusters based on the signal. The normalized average read count profile is plotted in the top panel, while the heatmaps in the bottom panel shows the normalized read count per enhancer where each row represents one enhancer annotated in the A549 cell line.

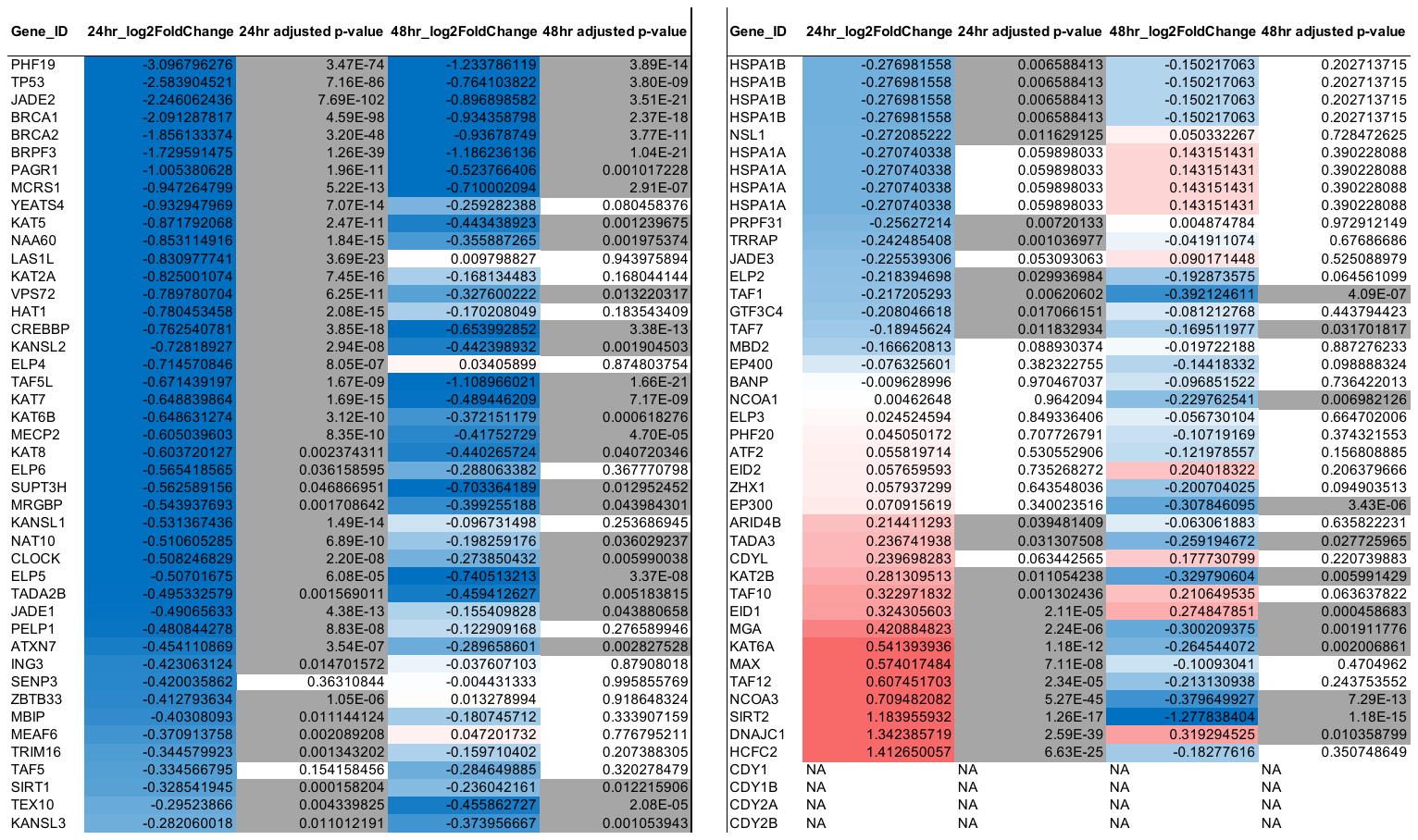

Table S 1:Differential expression of genes with histone acetyl transferase function. Log2 foldchange in expression from 24 hpt and 48 hpt cells are given in 2^nd^ and 4^th^ columns respectively. Their corresponding adjusted p-values are given in 3^rd^ and 5^th^ columns respectively. Log2 fold change in expression is color-coded from blue to red. Blue represents downregulated genes and red represents upregulated genes. Genes with significant expression changes with adjusted p-value < 0.5 are highlighted with grey color in the corresponding adjusted p-value column.
